# Two evolutionary histories in one nucleus: genome remodeling and allelic regulation underlying heterosis in hybrid oil palm

**DOI:** 10.64898/2026.08.27.747553

**Authors:** Xiangnian Su, Yanling Peng, Xuanwen Yang, Fan Zhang, Qi Xu, Zhiyao Ma, Yang Dong, Lianzhu Zhou, Hui Xue, Xuejing Cao, Zhi Zou, Yiwen Wang, Yongfeng Zhou, Xianhai Zeng

## Abstract

Oil palm (*Elaeis*) is the primary source of global vegetable oil. Interspecific hybrids of Elaeis exhibit pronounced heterosis by integrating two distinct subgenomes into a single nucleus, effectively combining the high yield of African oil palm (*E. guineensis*) with the high unsaturated fatty acid content and disease resistance of American oil palm (*E. oleifera*). However, the genetic basis underlying heterosis is still unclear. Here, we combine phased genome assembly, comparative genomics, evolutionary genomics and haplotype-aware transcriptomics to unravel the genetic architecture of heterosis of hybrid oil palm. We assemble the highly heterozygous F1 genome (’Reyou 40’, 3.75% heterozygosity) into a complete 1.73 Gb T2T haplotype (HapG) and a 1.84 Gb near-T2T haplotype (HapO with17 gaps). Despite 91.56% sequence identity, HapG and HapO diverged in LTR-RT occurrence and PAV affected genes, showing complementary biases in lipid metabolism and stress responses, respectively. Evolutionary genomics revealed that ancient WGDs preserved the palm family. Whereas lineage-specific lipid-related gene expansions in oil palm. Six ancient introgressed regions (∼64 Mb) in HapG were reshaped by transposable elements and tandem duplication, showing an enrichment of genes related to resistance and lipid metabolism. Transcriptomically, 82.2% of allelic gene pairs maintained balanced expression, accompanied by parental functional complementarity and dosage buffering, revealing a potential regulatory basis for coordinating parental genetic differences in the hybrid genome. These haplotype-resolved genomic resources offer vital targets for understanding heterosis and accelerating oil palm molecular breeding.

## Introduction

The palm family (*Arecaceae*) is a species-rich monocotyledonous lineage of major ecological and socioeconomic importance [1]. Among its members, the African oil palm (*Elaeis guineensis*) is the most productive oil crop per unit land area, supplying approximately 40% of global vegetable-oil demand while occupying less than 6% of the land devoted to oil crops [2, 3]. Its exceptional oil yield and agronomic performance have established *E. guineensis* as the predominant species underpinning the global palm-oil industry [1, 3, 4]. However, meeting rising vegetable-oil demand through continued plantation expansion would intensify pressure on tropical forests, carbon stocks and biodiversity [5–8]. Sustainable production must therefore rely increasingly on genetic improvement and climate-smart intensification to enhance yield, oil quality, disease resistance and resource-use efficiency [6, 9–11].

Although oil palm possesses outstanding oil production potential, its perennial nature, long generation cycle, and lengthy field evaluation period have greatly constrained the efficiency of conventional breeding [12, 13]. In contrast, the American oil palm (*E. oleifera*), native to Central and South America, represents an important wild genetic resource with valuable agronomic traits, including higher unsaturated fatty acid content [14, 15], reduced stem elongation [16], and enhanced resistance to multiple diseases [17]. These characteristics provide valuable genetic resources for oil palm improvement [18]. Therefore, by combining complementary parental alleles from *E. guineensis* and *E. oleifera*, interspecific hybridization and subsequent introgression provide a promising strategy to broaden the oil palm gene pool, pyramid favorable traits from divergent lineages, and develop cultivars with improved agronomic performance through heterosis [19–22]. However, despite the extensive application of *E. guineensis* × *E. oleifera* interspecific hybrids in oil palm breeding, the genomic basis underlying their reproductive compatibility remains largely unexplored. Successful interspecific hybridization requires the coordinated regulation of multiple reproductive processes, including pollen development, pollen–pistil interaction, pollen tube growth and guidance, and gamete fusion [23–25]. Although divergence between parental lineages can generate extensive genomic structural variation and sequence differentiation, the conservation of essential reproductive regulators may contribute to maintaining reproductive compatibility between divergent genomes [26, 27]. In other plant systems, genes involved in pollen formation, pollen recognition, and gamete fusion are often subjected to strong evolutionary constraints due to their indispensable roles in sexual reproduction [28–30]. However, whether such conserved reproductive modules are retained between *E. guineensis* and *E. oleifera*, and whether they contribute to successful interspecific hybridization, remain largely unknown.

Heterotic performance cannot be attributed solely to the presence of beneficial parental genes. Instead, the coexistence of two evolutionarily divergent chromosome sets forms a composite genome, where structural variations, repeat compositions, and gene content disparities collectively reconfigure gene regulatory networks [31–33]. Palm genomes retain an ancient whole-genome duplication shared by Arecaceae, after which subsequent chromosome rearrangements, gene loss and retention, and family turnover progressively reshaped modern genomes, driving karyotype diversification and the functional divergence of carbon and lipid metabolic genes [34–36]. Since their divergence, LTR-retrotransposon turnover, gene duplication, presence–absence variation and structural rearrangements may have progressively differentiated the genome architecture and gene repertoires of *E. guineensis* and *E. oleifera* [37, 38].

Historical gene flow between diverged lineages can generate introgressed genomic tracts whose ancestry differs from that of the surrounding chromosomal background [39, 40]. Once transferred, these tracts can be progressively fragmented and reshaped by recombination [39], while lineage-specific transposable element (TE) proliferation, TE-mediated structural rearrangements, and subsequent gene duplication or loss can further alter their sequence composition, gene content and functional potential [40–42]. In an F₁ nucleus, divergent parental chromosome complements share a common regulatory environment but can exhibit balanced, stably biased or developmentally switching allelic expression [43, 44]. These expression modes may maintain dosage-sensitive functions [45] and coordinate complementary parental contributions across development [45, 46].

Allelic expression differences can arise from cis-regulatory sequence divergence, copy-number variation, TE insertions and other structural rearrangements, whereas presence–absence variation (PAV) contributes to broader expression divergence at the gene-locus level [47–49]. Tandem duplication alters locus-level dosage through the combined output of multiple copies, while epigenetic and three-dimensional chromatin states further shape allelic accessibility and expression [50–52]. In an F1 hybrid, allelic expression differences within a shared trans environment indicate cis-regulatory divergence after controlling for mapping bias [31, 52]. Such effects have been documented in Arabidopsis, potato [53], cotton [54] and maize [51]. Genomic studies in *Elaeis* have identified candidate genes and regulatory pathways underlying mesocarp oil biosynthesis and accumulation, fruit development and agronomically important trait variation [3, 55–57]. Although chromosome-scale comparisons between *E. guineensis* and *E. oleifera* highlight overall genome conservation alongside local structural and repeat variations [38], current non-T2T reference genomes—derived from separate individuals—cannot directly resolve both parental haplotypes within a single hybrid genome. Single-parent references obscure haplotype-specific structural variants and bias allele-specific analyses [58]. In contrast, a haplotype-resolved hybrid genome resolves both parental chromosomes within a single native nucleus, enabling precise evaluation of haplotype dynamics and the genetic basis of heterosis in hybrid oil palm [59, 60].

In this study, we generated haplotype-resolved reference genomes for the interspecific F₁ hybrid oil palm ‘you 40’ comprising a T2T HapG genome and a near-T2T HapO genome. We systematically investigated haplotype divergence and genome evolution by integrating sequence variation, structural variation, transposable elements, gene content, the Dura genome, and representative Arecaceae species. Haplotype-resolved Hi-C maps and transcriptomes across five fruit developmental stages were further integrated to examine three-dimensional chromatin organization, allelic expression dynamics, and the transcriptional effects of tandem duplication. Specifically, we addressed four questions. First, how extensively have the two parental haplotypes diverged in sequence composition, repeat landscapes and gene content despite conserved chromosome-scale organization? Second, how have ancient introgression and subsequent TE- and duplication-associated remodeling shaped the defense and metabolic repertoire of HapG? Third, how have ancestral whole-genome duplication and lineage-specific gene-family turnover generated distinct lipid-metabolic and stress-response repertoires in the two parental lineages? Finally, how are these structural and functional differences coordinated through three-dimensional genome organization, allelic expression balance and tandem-copy regulation during fruit development? Together, this haplotype-resolved multi-omics framework links parental genome divergence, evolutionary remodeling and regulatory balance, providing a genomic basis for understanding heterosis and facilitating the breeding of high-yielding and resilient hybrid oil palm.

## Results

### Haplotype-specific PAVs show complementary functional specialization underlying heterosis

The American oil palm *(E. oleifera*) and African oil palm (*E. guineensis*) differ markedly in plant architecture, fruit morphology and agronomic traits (Figure 1A, B). HiFi *k*-mer analysis showed heterozygous and homozygous peaks at 49× and 98×, respectively, with an estimated heterozygosity of ∼3.75% (Figure S1A; Table S1, Supporting Information). The hybrid genome was assembled and phased using hifiasm [61], yielding two haplotype-resolved assemblies, HapG and HapO. Whole-genome *k*-mer phylogeny clustered HapG with *E. guineensis* and HapO with *E. oleifera*, confirming their respective parental origins (Figure S1B, Supporting Information). HapG yielded a complete 1.73-Gb T2T assembly comprising 16 gap-free chromosomes, whereas HapO yielded a 1.84-Gb near-T2T assembly comprising 16 chromosomes with 17 unresolved gaps. The contig N50 values of HapG and HapO were 123.55 Mb and 134.18 Mb, respectively, and their BUSCO [62] completeness reached 99.1% and 99.6% (Tables S2 and S3, Supporting Information). Protein-coding gene annotation identified 34,581 genes in HapG and 31,627 genes in HapO. Canonical plant telomeric repeats (CCCTAAA/TTTAGGG) were detected at chromosome ends. CENH3 ChIP-seq mapping delineated centromeres with diverse repeat architectures (Table S4, Supporting Information). The HapG chromosome 10 centromere lacked continuous arrays of canonical centromeric repeats and instead comprised an interspersed mosaic of Copia and Gypsy LTR-RTs, DNA/hAT transposons, rDNA repeats, and SSRs (Figure 1C).

**Figure 1.**
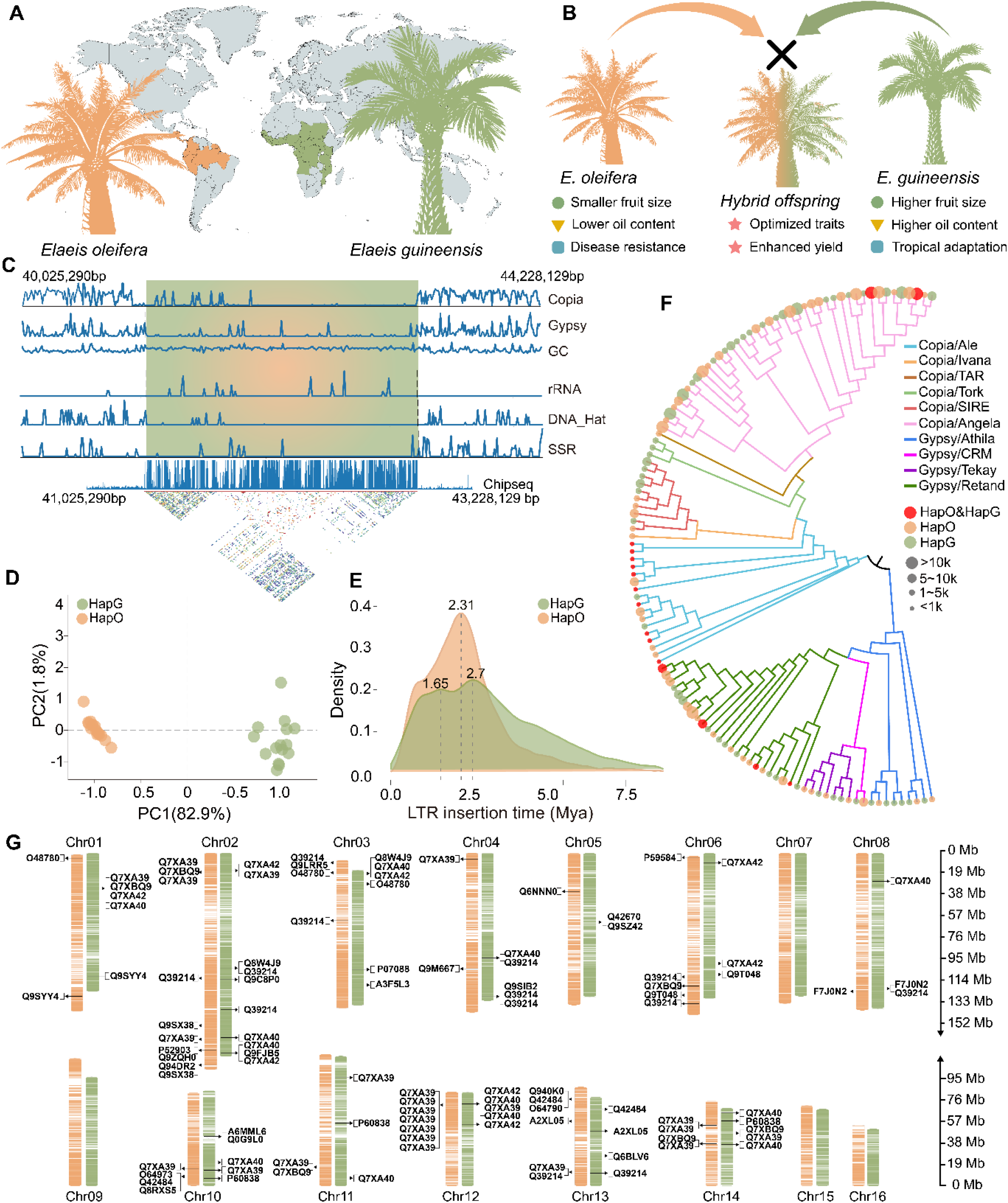
Haplotype-resolved assembly reveals divergent transposon landscapes and centromere architectures in an interspecific oil palm hybrid. (A) Geographic origins of *E. oleifera* and *E. guineensis*. (B) Morphological and agronomic characteristics of the parental species and their interspecific hybrid. (C) CENH3-defined centromeric region on HapG chromosome 10, showing repeat composition, GC content, CENH3 enrichment, and internal sequence similarity. (D) Principal component analysis of chromosomes based on haplotype-biased k-mers. (E) Insertion-time distributions and major amplification peaks of intact long terminal repeat retrotransposons (LTR-RTs). (F) Phylogeny of intact LTR-RTs inferred from conserved protein domains. Branch colors, tip colors and symbol sizes indicate LTR-RT lineages, haplotype origins and family abundance, respectively. (G) Chromosomal distribution of haplotype-specific presence–absence variation (PAV) genes in HapG and HapO, with genes related to stress resistance and lipid metabolism highlighted.

HapG and HapO had similar repeat content (77.54% versus 78.58%) and were both LTR-RT-dominated, but differed in Gypsy/Copia composition (56.9%/42.3% versus 51.5%/48.1%; Table S5; Figure S2A, B, Supporting Information). SubPhaser-based PCA [63] of haplotype-biased *k*-mers clearly separated HapG and HapO, consistent with their distinct repeat landscapes (Figure 1D). Although the insertion-time distributions of intact LTR-RTs broadly overlapped between the two haplotypes, their modal peaks differed markedly (Figure 1E). Phylogenetic analysis further revealed both shared and haplotype-specific LTR-RT families (Figure 1F). Thus, despite comparable overall repeat content, HapG and HapO exhibited distinct LTR-RT landscapes, reflecting differences in family composition and amplification history. We next examined haplotype-specific gene content. Despite broad sequence conservation between HapG and HapO (91.56% genome-wide similarity and 98.38% similarity across orthologous CDSs), the two haplotypes differed substantially in their gene complements (Figure S3A-C, Supporting Information). We identified 1,808 HapG-specific and 1,285 HapO-specific PAV genes. Of these, 465 HapG-specific and 340 HapO-specific genes were organized into 82 and 85 genomic clusters, averaging 5.7 and 4.0 genes per cluster, respectively (Figure 1G; Tables S6 and S7). Representative HapG clusters contained GDSL lipase genes and genes involved in organellar energy metabolism, whereas HapO clusters contained NB-ARC/LRR-type disease-resistance genes (Figure S4A-C, Supporting Information). Functional annotation identified 104 and 26 oil-related PAV genes in HapG and HapO, respectively, together with 108 and 137 PAV genes associated with environmental responses (Figure S4D and Figure S16, Supporting Information). Nearly all PAV genes were TE-associated, either without accompanying SVs (82.4% in HapG and 72.7% in HapO) or together with SVs (17.1% and 26.6%, respectively; Figure S5, Supporting Information).

### Comparative genomics of the palm family reveals lipid gene expansion in oil palm

We first performed MCScanX-based [64] synteny analysis of HapG, HapO, *Cocos nucifera*, *Areca catechu*, Phoenix *dactylifera*, *P. roebelenii*, and *Asparagus officinalis*. HapG and HapO exhibited extensive conservation of chromosome structure and gene order, whereas the extent of collinearity declined with increasing phylogenetic distance (Figure 2A; Table S8). Complementary OrthoFinder analysis [65] assigned 184,602 predicted genes (89.7%) to 21,932 orthogroups, including 7,651 orthogroups shared across all seven assemblies and 1,626 single-copy orthogroups (Figure S6A; Table S9). Gene-family presence–absence analysis grouped HapG and HapO together and identified 373, 181, and 128 families specific to HapG, HapO, and coconut, respectively (Figure S6B). In addition, 1,042 orthogroups were shared exclusively between HapG and HapO (Figure 2B; Table S9). Functional enrichment analysis showed that HapG-specific genes were primarily associated with ribosome biogenesis, translation and defense-related processes, whereas HapO-specific genes were associated with hormone biosynthesis, kinase signaling and environmental responses (Figure S6C, D).

**Figure 2.**
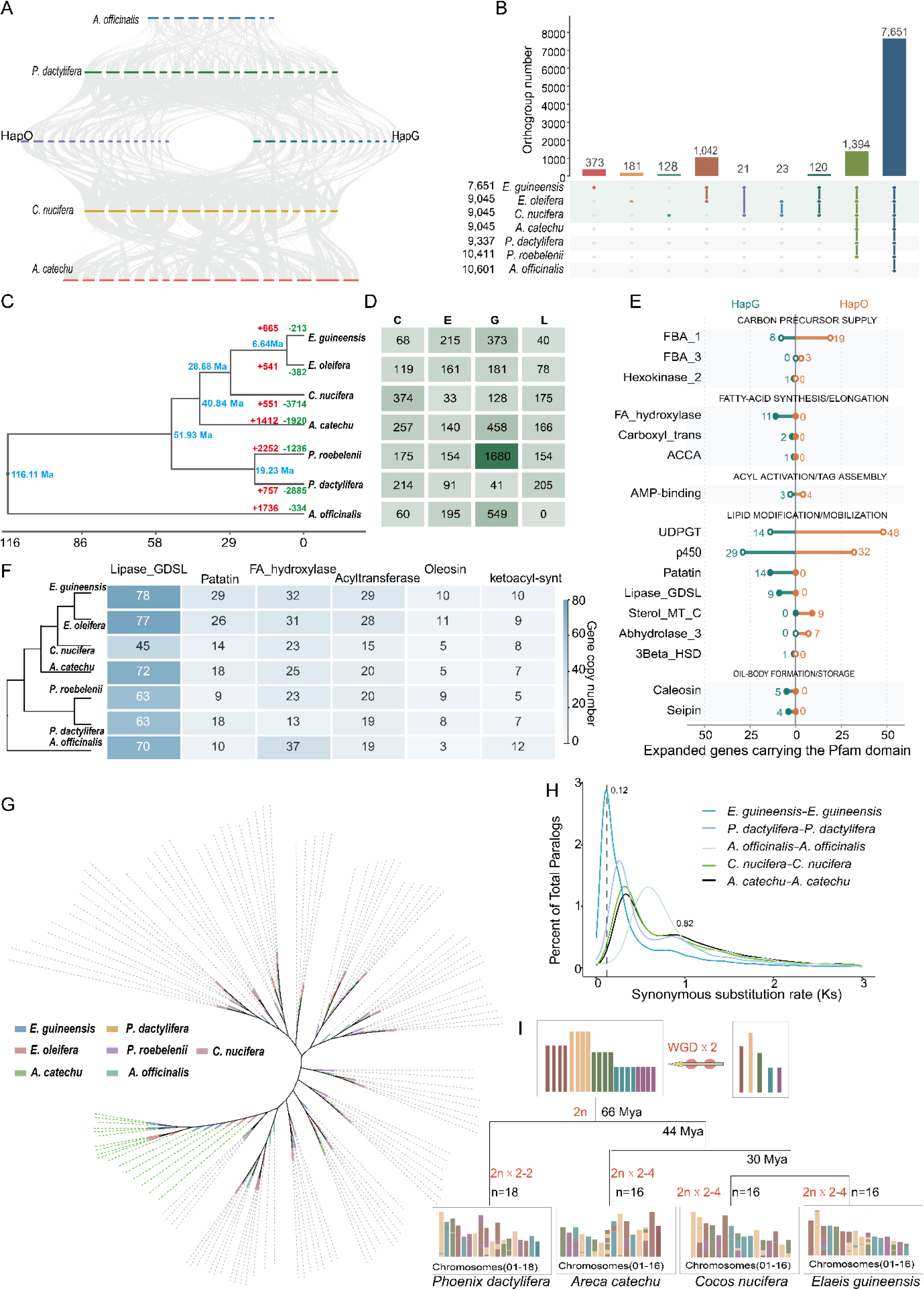
Ancestral genome duplication and lineage-specific expansion of lipid-related gene families in oil palm. (A) Chromosome-scale synteny among representative palms and *Asparagus officinalis*. (B) Shared and species-specific orthogroups across the examined species. (C) Species phylogeny, divergence times, and gene-family expansion and contraction. Red and green indicate significantly expanded and contracted gene families, respectively, and blue indicates divergence times. (D) Gene-family turnover across the examined species. The matrix summarizes the numbers of contracted (C), expanded (E), gained (G), and lost (L) gene families. (E) Functional classification of expanded genes in HapG and HapO. Solid and open circles denote direct functional associations and broad or context-dependent domain associations, respectively. (F) Copy-number variation of lipid-related Pfam families across species. (G) Phylogeny of Patatin domain-containing genes across representative palms and *Asparagus officinalis*; colors indicate species origins. (H) *Ks* distributions of syntenic paralogs, with major peaks indicated. (I) Ancestral karyotype reconstruction across representative palm lineages. Colors denote ancestral chromosomal components, and circles indicate whole-genome duplication (WGD) events.

The time-calibrated phylogeny estimated that the parental lineages represented by HapG (*E. guineensis*) and HapO (*E. oleifera*) diverged approximately 6.64 Ma and split from the *C. nucifera* lineage approximately 28.68 Ma. Following species divergence, the two oil palm lineages underwent markedly asymmetric gene-family expansion and contraction: 665 families expanded and 213 contracted in *E. guineensis*, whereas 541 expanded and 382 contracted in *E. oleifera* (Figure 2C, D; Table S10). Expanded gene families exhibited clear lineage biases (Figure S7A, B, Supporting Information). Expanded genes in *E. guineensis* were primarily associated with fatty-acid synthesis and elongation, lipid mobilization, oil-body formation and defense signaling, whereas those in *E. oleifera* were associated with carbon-precursor supply, lipid modification, immune recognition, abiotic-stress responses and redox detoxification (Figure 2E; Figure S7C, D, Supporting Information). Comparison of six lipid-related gene families showed that the two oil palm genomes generally contained more genes involved in lipid remodeling, fatty-acid metabolism and oil-body formation than the other palm genomes examined (Figure 2F; Table S11, Supporting Information). The Patatin family formed multiple expanded clades in *Elaeis* and showed differential copy-number retention between HapG and HapO (Figure 2G). The *Ks* distribution of syntenic paralogues showed two distinct peaks at approximately 0.82 and 0.12 (Figure 2H), corresponding to two ancient whole-genome duplication (WGD) events. WGDI-based [66] ancestral karyotype reconstruction revealed extensive retention of ancestral chromosomal segments across Arecaceae, together with lineage-specific chromosomal fusions, fissions, and translocations; *Elaeis* and coconut retained particularly similar chromosomal compositions (Figure 2I).

### Ancient introgression of resistance genes in hybrid oil palm

Despite extensive collinearity across all 16 homologous chromosome pairs, sliding-window analysis revealed pronounced regional heterogeneity in nucleotide divergence (π) between HapG and HapO (Figure 3A). Genome-wide comparisons incorporating the newly assembled African oil palm Dura genome further showed extensive chromosome-scale synteny, but greater structural divergence between HapG and HapO than between Dura and HapG, as reflected by the increased number and cumulative span of structural variants (Figure S9). Notably, several key reproduction-associated loci remained highly conserved despite this genome-wide structural divergence. The CYP703A2-like–GPAT region, PRK-family locus, and HAP2/GCS1-like locus all showed strong conservation of genomic organization and/or protein sequence between HapG and HapO (Figures S10A-D, Supporting Information).

**Figure 3.**
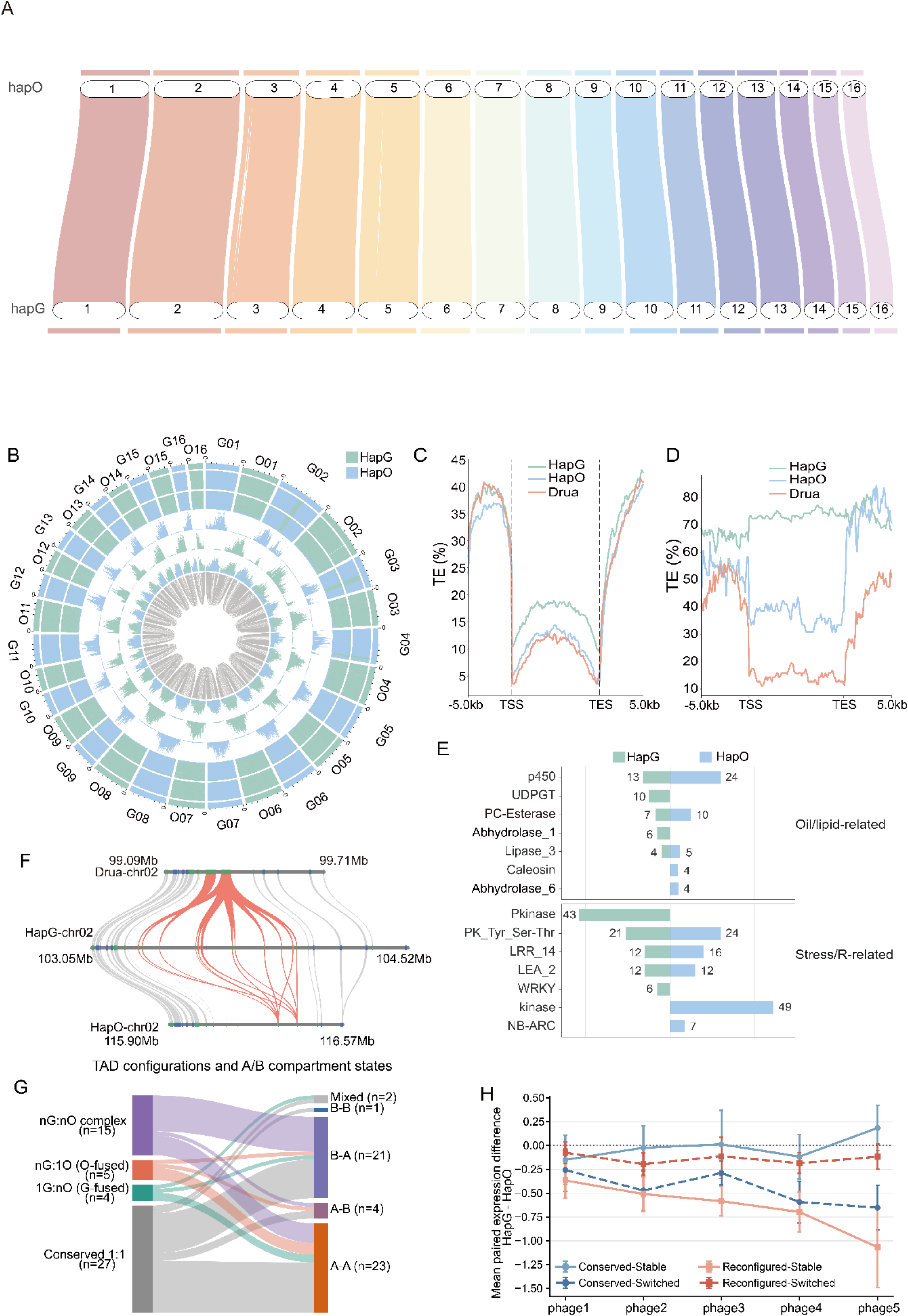
Evolutionary and regulatory features of ancient introgressed regions in HapG. (A) Chromosome-scale synteny and sequence divergence between HapG and HapO. (B) Genomic features, collinear relationships, and candidate introgressed regions in HapG and HapO. (C, D) TE occupancy across all genes (C) and PAV genes (D), including their 5-kb flanking regions, within the HapG introgressed regions and corresponding homologous intervals in HapO and Dura. Dashed lines indicate transcription start sites (TSSs) and transcription end sites (TESs). (E) Copy-number comparison of selected oil-related and stress-related gene families within the HapG introgressed regions and corresponding HapO intervals. (F) Local synteny of the RPM1 gene cluster across Dura, HapG, and HapO on chromosome 2. (G) TAD configurations and A/B compartment states of paired introgressed intervals. (H) Developmental expression profiles of HapG genes grouped by TAD configuration and compartment state. Data are presented as means ± s.e.m.

SubPhaser [63] identified six candidate ancient introgressed regions enriched in HapO-specific k-mers on HapG chromosomes Chr02, Chr03, Chr04, Chr06, Chr09 and Chr15. These regions collectively spanned approximately 64 Mb, accounting for 3.70% of the HapG genome (Figure 3B; Table S12, Supporting Information). Comparisons of homologous intervals among HapG, HapO and the newly assembled *E. guineensis* Dura genome showed that all six HapG regions were more closely related to their HapO counterparts than to Dura (mean p-distance for HapG–HapO versus HapG–Dura: 0.0164 versus 0.0249). Their inferred age of approximately 4.0 Ma was younger than the estimated 6.64-Ma divergence between *E. guineensis* and *E. oleifera*, supporting their identification as ancient introgressed regions in HapG (Figures S11 and S12A, Supporting Information). The introgressed cores exhibited significantly lower π values than their ±3-Mb flanking intervals, and genes within these regions had lower Ka/Ks ratios than background genes (0.395 versus 0.433; P < 0.001), indicating reduced nucleotide diversity (Figure S12B, C). The six ancient introgressed regions in HapG differed from their homologous intervals in HapO and Dura in gene content, tandem duplication and repeat composition (Figure S12D and Table S13, Supporting Information). Gypsy and Copia elements were expanded within the HapG introgressed regions, which showed higher gene-associated TE coverage than the corresponding HapO and Dura intervals. TE coverage approached 70% across the transcribed regions of HapG-specific PAV genes (Figure 3C, D), revealing extensive TE-associated local genome remodeling. Resistance-gene expansion was the most prominent functional feature of these regions. Phylogenetic analysis across Arecaceae revealed expansions of multiple resistance-gene lineages in oil palm (Figures S13, S13B, Supporting Information; Table S12, Supporting Information). Local synteny analysis further localized an expanded RPM1 tandem cluster to the ancient introgressed region on HapG Chr02 (Figure 3F), revealing an earlier expansion in the oil palm lineage followed by more recent local amplification in HapG. The HapG introgressed regions also contained more UDP-glycosyltransferase and WRKY family members, whereas their HapO homologues contained more caleosin, cytochrome P450 and NB-ARC genes (Figure 3E). Highly differentiated segments and PAV genes in HapG were primarily associated with immune and stress responses, lipid metabolism, and energy regulation (Figures S14A, B; Table S14, Supporting Information).

Hi-C comparison of the six ancient introgressed regions in HapG and their homologous regions in HapO identified 51 paired TAD events, comprising 27 conserved 1:1 events and 24 reconfigured events, including 15 complex nG:nO, five nG:1O (HapO-fused), and four 1G:nO (HapG-fused) events (Figure 3G and Table S15, Supporting Information). Regional reconfiguration rates ranged from 33.3% to 60.0%. Compartment correspondence patterns were region-specific, comprising A-A on chr02, B-A on chr03/04/15, A-B on chr09, and B-B or mixed states on chr06 (Figure S15A, Supporting Information).

Within HapG, genes in A compartment showed significantly higher expression than those in B compartments (P = 1.8 × 10^-5^ Figure 3H). All six homologous regions showed higher median expression in HapO than in HapG (regional sign-flip test, P = 0.03125) (Figure S15B, Supporting Information); median log^2^expression was 3.021 in HapG and 3.195 in HapO, with a median paired HapG−HapO difference of −0.259, and 41 of the 51 events showed higher expression in HapO (Figure S15C, Supporting Information). No significant differences were detected in the overall reconfiguration rate, TAD length, or compartment-switching frequency, and expression differences were associated with neither TAD reconfiguration nor compartment switching (all P > 0.05; Figure S15D).

### Allelic Expression Balance and Tandem Duplication Coordinate Complementary Parental Contributions to Heterosis in Hybrid Oil Palm

We identified 26,034 allelic gene pairs, together with 8,547 HapG-specific and 5,593 HapO-specific genes (Figure 4A). Based on coding-sequence (CDS) conservation, allelic pairs and haplotype-specific genes were classified as CDS-identical, CDS-diverged, or haplotype-specific. TE coverage progressively increased across these categories, from CDS-identical to CDS-diverged and haplotype-specific genes (Figure 4B, C), suggesting that increasing sequence divergence is associated with increasingly TE-rich genomic environments.

**Figure 4.**
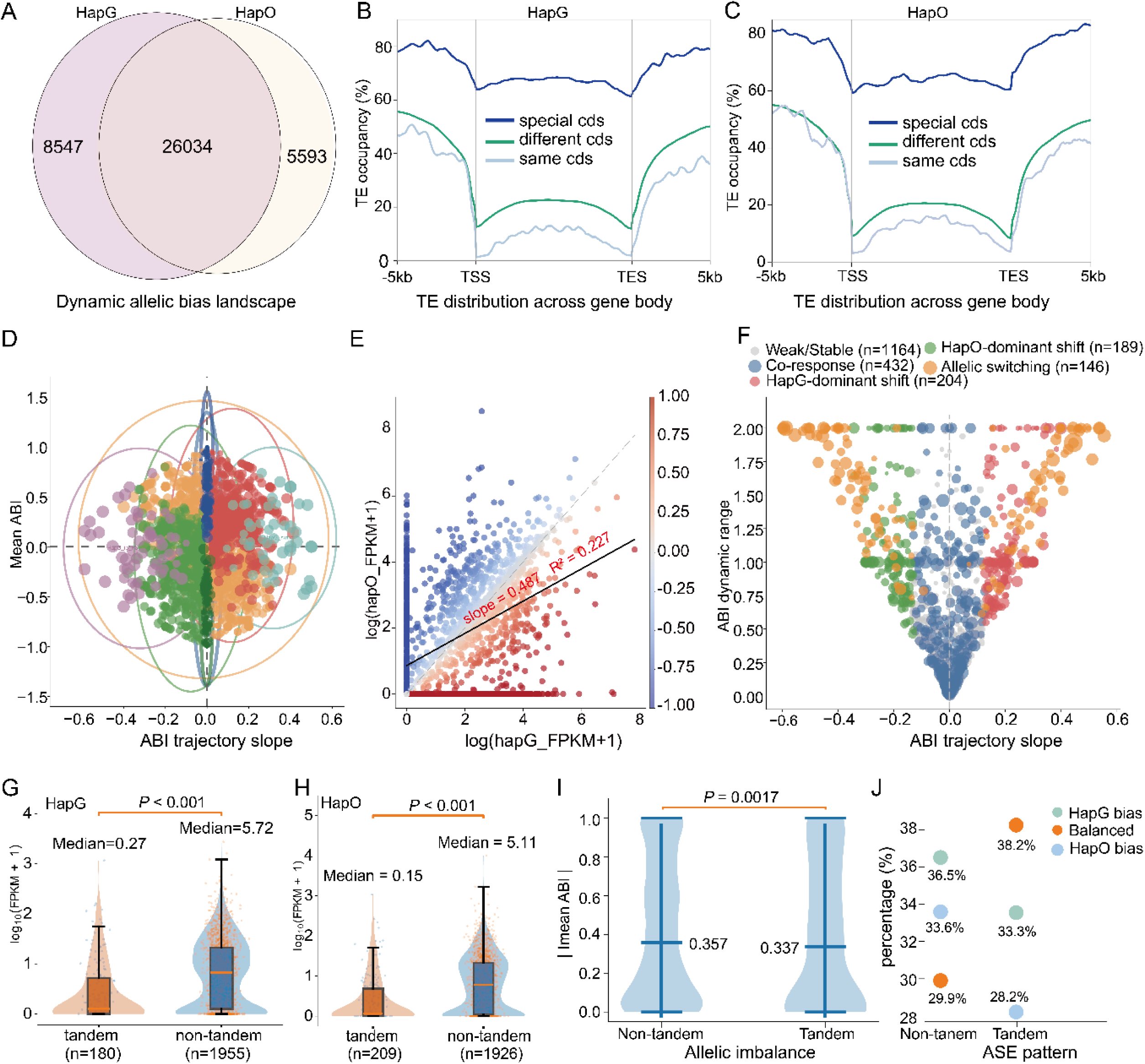
Allelic expression dynamics and duplication-associated dosage regulation in hybrid oil palm. **(A)** Shared and haplotype-specific genes between HapG and HapO. (B, C) TE occupancy across HapG (B) and HapO (C) genes and their ±5-kb flanking regions, grouped as CDS-identical, CDS-diverged, and haplotype-specific. TSS and TES indicate transcription start and end sites, respectively. (D) Allelic-bias landscape of 21,178 expressed allelic pairs. Each bubble represents one allelic pair, with ABI trajectory slope on the x-axis and mean ABI across five developmental stages on the y-axis. Bubble size indicates ABI dynamic range, and colors denote eight expression classes: balanced (n = 14,424), dynamically balanced (n = 2,994), HapG-dominant shift (n = 1,228), HapO-dominant shift (n = 1,461), stable HapG bias (n = 673), stable HapO bias (n = 293), HapG-to-HapO switch (n = 57), and HapO-to-HapG switch (n = 48). Positive and negative mean ABI values indicate HapG and HapO bias, respectively; positive and negative slopes indicate developmental shifts toward HapG and HapO. Colored ellipses summarize the distribution of each class. (E) Relationship between HapG and HapO expression across allelic pairs. Point color represents mean ABI. The solid line shows the linear fit, ln(HapO + 1) = 0.487 × ln(HapG + 1) + 0.864 (R² = 0.227), and the dashed line indicates y = x. (F) Dynamic allelic-bias patterns of 2,135 allelic pairs. Bubble size represents ABI dynamic range, and colors indicate weak/stable (n = 1,164), co-response (n = 432), HapG-dominant shift (n = 204), HapO-dominant shift (n = 189), and allelic switching (n = 146). (G, H) Per-copy expression of tandem and non-tandem genes in HapG (G) and HapO (H). Median values and significance are indicated in the plots. (I) Allelic imbalance of tandem and non-tandem loci based on |mean ABI|; tandem loci showed lower imbalance (0.337 vs. 0.357; P = 0.0017). (J) Proportions of HapG-biased, balanced, and HapO-biased loci in tandem and non-tandem groups; percentages are indicated in the plot.

We next examined whether genomic divergence between the two parental haplotypes was accompanied by transcriptional divergence. Among the 21,178 allelic gene pairs retained after expression filtering, 82.2% exhibited balanced expression, including 14,424 consistently balanced and 2,994 dynamically balanced pairs. The remaining 3,760 pairs showed non-balanced expression, comprising 1,228 and 1,461 pairs with dynamic shifts toward HapG- and HapO-biased expression, respectively, 673 and 293 pairs with persistent HapG and HapO bias, respectively, and 105 pairs exhibiting complete reversals in bias direction. Functional enrichment revealed distinct haplotype-associated profiles, with HapG-biased genes predominantly enriched in defense responses and lipid biosynthesis, whereas HapO-biased genes were associated with hormone-mediated stress responses and lipid transport and modification (Figures 4D and S16A, B; Table S16).

Despite the predominance of balanced expression, expression levels between allelic pairs were only weakly correlated (slope = 0.487, R² = 0.227; Figure 4E), indicating substantial quantitative variation between parental alleles. Across five fruit developmental stages, 2,135 dynamically regulated allelic pairs were further resolved into distinct temporal expression patterns, including weakly biased/stable, biparentally co-responsive, and directionally shifted/switching patterns (Figure 4F; Figure S17A). Stage-specific enrichment further revealed functional partitioning between the two haplotypes: HapG-biased genes were preferentially associated with carbohydrate phosphorylation, galactokinase activity, fatty acid biosynthesis, MAPK signaling, and drought responses, whereas HapO-biased genes were preferentially associated with fatty acid transport, glycerol-3-phosphate acyltransferase activity, and hormone-mediated stress responses (Figures S17B and S17C, Table S17).

Tandem copy expansion was not accompanied by a proportional increase in transcriptional output. Per-copy expression of tandemly duplicated genes was markedly lower than that of non-tandem genes in both HapG (median FPKM, 0.27 vs. 5.72) and HapO (0.15 vs. 5.11; *P* < 0.001; Figure 4G, H; Table S18, Supporting Information). Tandem loci also showed a lower mean absolute allelic-bias index than non-tandem loci (|ABI|, 0.337 vs. 0.357; *P* = 0.0017; Figure 4I). Balanced expression was more frequent among tandem loci than among non-tandem loci (38.2% vs. 29.9%), whereas the proportions of HapG- and HapO-biased loci were correspondingly lower (Figure 4J; Table S19, Supporting Information). Overall, tandem loci were characterized by lower per-copy expression, reduced allelic-bias magnitude, and a higher proportion of balanced expression.

## Discussion

In this study, we investigated the genetic basis of heterosis in hybrid oil palm by integrating phased genome assembly, comparative genomics, evolutionary genomics and haplotype-aware transcriptomics: (1) HapG and HapO retained extensive chromosome-scale synteny but shared only 91.56% genome-wide sequence identity, indicating substantial sequence divergence between the parental haplotypes. The PAV-related genes in the two haplotypes showed functional differentiation associated with lipid metabolism and environmental responses. (2) Comparative genomics across the palm family revealed a broadly conserved chromosomal evolutionary framework and shared ancestral genome-duplication signals, whereas the two parental oil palm lineages underwent distinct patterns of gene-family expansion and contraction after divergence. (3) Ancient introgressed regions in HapG showed local expansion of resistance genes, including RPM1. (4) Fruit transcriptomic analyses revealed a coordinated regulatory pattern characterized by predominantly balanced allelic expression, complementary parental functional biases, and patterns consistent with dosage buffering.

### TE dynamics and ancient introgression shape multilayered haplotype divergence

HapG and HapO contained similar overall proportions of repetitive sequences but differed markedly in LTR-RT composition, insertion-time peaks and haplotype-restricted lineages, while most haplotype-specific PAV genes occurred in TE-associated regions. These results identify TE dynamics as an important component of haplotype divergence [67–69], paralleling lineage-specific TE turnover in other diploid and polyploid plants [70–72]. Shared and haplotype-restricted LTR-RT clades are consistent with ancestral element retention followed by asynchronous amplification and differential loss after the divergence of *E. guineensis* and *E. oleifera*. Similar repeat abundance therefore masks distinct histories that established different structural and gene-content backgrounds in the parental haplotypes. Therefore, TE-driven genome remodeling has shaped the divergent genomic backgrounds of the two parental subgenomes, HapG and HapO, whereas the conservation of reproductive-associated gene modules reflects the stable maintenance of functional core regions during hybrid formation. This combination of ‘structural divergence and functional conservation’ may represent an important molecular basis underlying the success of interspecific hybridization.

Six HapG regions were more similar to HapO than to the newly assembled *E. guineensis* Dura genome, and their inferred ages postdated the divergence of the two oil palm species. Together with reduced nucleotide diversity and lower Ka/Ks ratios, these patterns define ancient introgressed tracts in the African oil palm-derived haplotype. Introgressed segments can persist locally without erasing recipient-genome identity, as reported in date palm and other perennial crops [73–75]. Enrichment of immune, stress-response, lipid-metabolic and energy-regulatory functions suggests that ancient gene flow broadened the repertoire inherited through HapG. However, enrichment and retention do not establish adaptive introgression; population-level selection tests and functional and phenotypic validation remain necessary [76, 77]. These tracts predate the present F₁ hybrid and represent components of the HapG parental legacy rather than products of recent hybridization.

The HapG introgressed regions showed increased Gypsy and Copia abundance, elevated gene-associated TE coverage and extensive TE occupancy across HapG-specific PAV genes, together with local resistance-gene duplication. Thus, the introgressed segments were subsequently remodeled rather than retained as static blocks, consistent with the evolutionary lability of introgressed DNA in plants [69, 78, 79] and defense-related post-introgression variation in poplar and rice [69, 79, 80]. The clearest example is the expanded *RPM1* cluster on chromosome 3: R-gene phylogenies indicate an earlier oil palm expansion followed by recent tandem duplication within HapG. This pattern accords with rapid NLR evolution through duplication, recombination and TE-associated rearrangement [81–83]. Although Arabidopsis *RPM1* encodes an intracellular immune receptor [84], its expansion in HapG indicates defense-related potential rather than direct control of bud rot resistance. Field variation among oil palm accessions and OxG progenies requires population and phenotypic validation [85, 86]. The defined intervals provide targets for testing copy-number, expression and trait associations of *RPM1* and neighboring defense loci.

Hi-C analysis identified conserved and reconfigured TADs across the introgressed regions, but TAD reconfiguration was not coupled to compartment switching, and B-to-A switching in HapG did not produce uniform HapG-biased activation. Expression across homologous intervals was instead slightly higher in HapO. This uncoupling argues against a model in which introgression generally opens chromatin and activates transcription, despite chromatin reorganization reported in plant hybrids and polyploids [87–90]. The tracts appear to have been accommodated within HapG chromatin in a locus-specific manner, with compartment state acting alongside local sequence and cis-regulatory differences. Because TEs can promote heterochromatin and alter nearby regulatory landscapes [32, 91], their accumulation may contribute to this variability. Three-dimensional architecture therefore modulates, but does not determine, the regulatory consequences of introgression.

### Genomic and regulatory foundations of heterosis in hybrid oil palm

The conserved chromosomal framework and shared ancestral WGD history of the two oil palm lineages contrast with their asymmetric gene-family turnover after divergence. This lineage-specific remodeling preferentially reinforced fatty-acid synthesis, lipid mobilization and oil-body formation in HapG, while enriching lipid modification, immune recognition and abiotic-stress responses in HapO. Ancestral WGD may therefore have supplied a common pool of duplicated genes [92–94], which was subsequently reshaped through differential retention, loss and small-scale duplication [92, 93, 95]. The resulting functional differentiation provides a plausible basis for parental complementarity in the hybrid, potentially integrating the lipid-production capacity of HapG with the metabolic flexibility and stress-response potential of HapO.

Among expressed allelic pairs, 82.2% maintained balanced expression, while a smaller fraction showed persistent, developmental or switching biases toward either parental haplotype. HapG- and HapO-biased genes were enriched in different lipid-metabolic and stress-response functions, indicating localized deployment of parental alleles without genome-wide dominance. Because both alleles experience the same trans-regulatory environment, this combination of broad balance and local bias is consistent with shared trans regulation attenuating some parental differences while stronger cis-regulatory divergence remains detectable [31, 43, 96–98]. Comparable patterns in maize [99, 100], rice and Arabidopsis [31] show that promoter variation, trans-acting factors and compensatory cis-trans interactions can jointly stabilize hybrid expression [101]. Such regulation may preserve parental functional diversity while maintaining transcriptional homeostasis across core processes [43, 102, 103]. Direct parental transcriptomes will nevertheless be required to distinguish hybrid-specific buffering from inherited parental expression differences.

Tandemly duplicated genes showed substantially lower expression per copy than non-tandem genes, together with a modestly lower absolute allelic-bias index and a higher proportion of balanced loci. Thus, tandem copy-number expansion was not translated into proportional transcriptional output and was associated with reduced allelic imbalance. Shared local chromatin or regulatory constraints, as well as regulatory partitioning among duplicates, could produce such nonlinear dosage responses [104, 105], but the present data cannot distinguish these mechanisms [106]. The reduced imbalance at tandem loci is compatible with partial buffering of haplotype copy-number asymmetry in the hybrid environment [107], whereas bias at non-tandem loci may more directly reflect haplotype-specific cis regulation [108].

Collectively, the data support a model in which lineage-specific TE turnover, ancient introgression, gene-family diversification and local duplication generated complementary parental genome backgrounds, while broad allelic balance and locus-specific regulation accommodated their coexistence in the F₁ hybrid. Heterosis in ‘Reyou 40’ is therefore more plausibly associated with the integration of complementary gene repertoires and regulatory stability than with uniform activation of one parental genome or a single introgressed locus. This framework identifies testable genomic regions and regulatory processes for population-level validation and haplotype-informed oil palm breeding.

## Materials and Methods

### Plant materials and sequencing

An F₁ hybrid between *E. guineensis* and *E. oleifera* (‘Reyou 40’), developed by the Chinese Academy of Tropical Agricultural Sciences, was used in this study. Fresh young leaves were collected and immediately frozen in liquid nitrogen. High-molecular-weight DNA was extracted using the cetyltrimethylammonium bromide method and purified with a Qiagen Genomic Kit. DNA quantity and purity were assessed using Qubit and NanoDrop instruments, and integrity was examined by agarose-gel electrophoresis. PacBio HiFi and Oxford Nanopore ultra-long libraries were sequenced on the PacBio Sequel II and PromethION platforms, respectively. Hi-C libraries were prepared from young leaves using proximity ligation and sequenced on an Illumina NovaSeq platform.

### Genome Assembly and Quality Evaluation

#### Genome survey

Genome size, heterozygosity and repetitive-sequence content were estimated from the k-mer frequency distribution of PacBio HiFi reads using GenomeScope 2.0 [109].

#### Haplotype-resolved genome assembly

Haplotype-resolved assembly was performed using hifiasm [61] by integrating PacBio HiFi, Oxford Nanopore ultra-long and Hi-C sequencing data. The two parental haplotypes were assembled independently. Contigs were grouped, ordered, oriented and anchored into chromosome-scale scaffolds using Juicer (v1.5) [110] and 3D-DNA (v180922) [111], followed by manual inspection and correction of Hi-C contact maps in Juicebox [112]. Remaining assembly gaps were examined using long-read alignments generated with minimap2 [113] and sequence-similarity searches performed with BLAST [114].

#### Assembly quality assessment

Gene-space completeness was evaluated using BUSCO [62] against the embryophyta_odb10 dataset. Assembly contiguity was evaluated using QUAST (v5.2.0) [115], which was used to calculate contig N50 and chromosome-level assembly statistics.

### Identification of telomeres and centromeres

#### Telomere identification

Canonical plant telomeric repeats, represented by the motif (TTTAGGG)n and its reverse complement (CCCTAAA)n, were identified using tidk [116]. The distribution of these repeat arrays at chromosome termini was used to evaluate telomere completeness.

#### Centromere identification

CENH3 ChIP-seq and matched input reads were quality-filtered using fastp (v0.23.2) [117] and aligned to the corresponding haplotype assemblies with Bowtie2 (v2.5.1) [118] using the parameters --very-sensitive --no-mixed --no-discordant -k 200 --maxins 800. Alignments with mapping quality below 2 were discarded, and PCR duplicates were removed using SAMtools (v1.10) [119]. CENH3 enrichment was calculated with bamCompare in deepTools [120] (v3.5.1) as log^2^ (ChIP/Input) coverage in 50-bp bins. Broad CENH3-enriched domains were identified using MACS2 [121] with the parameters --nomodel --extsize 291 --keep-dup auto --broad and visually inspected in IGV [122]. Continuous genomic regions consistently supported by the CENH3 enrichment profiles and broad-peak signals were defined as candidate centromeric regions.

### Gene and repeat annotation

Repetitive elements were annotated independently for HapG and HapO using complementary de novo, homology-based and structure-based approaches. Haplotype-specific repeat libraries were constructed with RepeatModeler [123] (v2.0.4), incorporating RepeatScout [124] (v1.0.6) and RECON [125] (v1.08). Intact LTR retrotransposons (LTR-RTs) were identified using LTR_FINDER (v1.07) [126] and LTRharvest [127] implemented in GenomeTools [128] (v1.6.1), and the resulting candidates were integrated and refined with LTR_retriever (v2.9.0) [129]. Redundant repeat consensus sequences were removed using CD-HIT (v4.8.1) [130], and tandem repeats were identified with Tandem Repeats Finder [131] (v4.09). Genome-wide repeat annotation was then performed with RepeatMasker [132] using the haplotype-specific libraries. Elements remaining unclassified were assigned with DeepTE [133], whereas intact LTR-RTs were further classified with TEsorter [134] according to their conserved protein domains. The resulting annotations were merged to generate repeat-masked HapG and HapO assemblies for gene prediction.

Protein-coding genes were predicted independently for HapG and HapO using the repeat-masked chromosome assemblies. Gene models were constructed by integrating ab initio predictions, transcript-derived evidence and homologous proteins from related plant species. Transcript evidence was used to define splice junctions and exon–intron boundaries, whereas protein alignments were used to support coding regions and refine gene structures. The three evidence sources were integrated to generate non-redundant consensus gene models. Low-confidence models with extensive overlap with annotated transposable elements and no transcript or protein-homology support were excluded. Coding sequences from the retained gene models were translated to generate the predicted proteomes of HapG and HapO. Conserved protein domains were identified by searching the predicted proteins against Pfam-A using hmmscan [135], with domain matches retained according to the curated gathering threshold of each Pfam family.

### Haplotype comparison, PAVs and LTR-retrotransposon evolution

#### Chromosome synteny and structural variation

GENESPACE [136] was used to identify chromosome-scale synteny, collinear gene blocks and homologous chromosome correspondence between HapG and HapO. Whole-genome alignments were generated with MUMmer [137], and structural variants were identified using SyRI [138] and visualized with plotsr [139]. Pairwise nucleotide divergence was calculated in sliding windows across callable whole-genome alignments.

#### Presence–absence variation

Haplotype-specific presence–absence variation (PAV) genes were identified according to homologous gene relationships and conserved syntenic positions between HapG and HapO. Spatially adjacent PAV genes were further grouped into local genomic blocks based on their chromosomal proximity.

#### Haplotype-specific k-mer analysis

SubPhaser [63] was used to identify haplotype-biased 15-mers, assign chromosomes to their respective parental origins and characterize local variation in haplotype-specific k-mer abundance across the two chromosome complements.

#### LTR-retrotransposon evolution

Conserved protein domains of intact LTR retrotransposons were identified using TEsorter. Domain sequences longer than 100 amino acids were retained and aligned for phylogenetic inference. The insertion time of each intact LTR-RT was estimated from the sequence divergence between its paired terminal repeats.

### Identification and evolutionary characterization of candidate introgressed regions

#### Identification and analysis of candidate introgressed regions

Contiguous HapG regions showing enrichment of HapO-biased k-mers relative to the surrounding HapG background were defined as candidate introgressed intervals. After excluding indels, missing sites and low-quality variants, consensus sequences were reconstructed from high-quality biallelic SNPs. Pairwise nucleotide divergence was calculated as the number of nucleotide differences divided by the callable alignment length, and divergence time was estimated as T = d/(2μ), where d denotes pairwise nucleotide divergence and μ is the nucleotide substitution rate.

#### Validation of introgressed ancestry

For independent validation, the six candidate HapG intervals were aligned against an E. guineensis Dura genome using nucmer [137] with the parameters --maxmatch -l 100 -c 500. One-to-one alignments were retained using delta-filter -1, and overlapping alignment coordinates were merged to define the corresponding homologous intervals in Dura. Pairwise p-distances were then calculated among the homologous HapG, HapO and Dura sequences to evaluate their relative sequence affinities.

### Identification of intrachromosomal collinear copies

Nucleotide-level collinearity between the candidate HapG intervals and their homologous Dura regions was determined from MUMmer whole-sequence alignments. Protein-level collinearity was subsequently assessed using WGDI [66], based on homologous protein pairs and their genomic coordinates, to identify intrachromosomal collinear blocks and potential gene duplications within HapG.

### Comparative phylogenomics, gene-family evolution and ancestral karyotype reconstruction

#### Orthogroup inference and phylogenomic analysis

Using MCScanX [64], chromosome-scale collinearity was evaluated among HapG, HapO, C. nucifera, A. catechu, P. dactylifera, and A. officinalis. Orthogroups and single-copy orthologs across these species and P. roebelenii were subsequently identified using OrthoFinder [65]. P. roebelenii was omitted from the synteny analysis because its assembly lacks chromosome-level anchoring, but it was included in the OrthoFinder analysis for orthogroup and single-copy ortholog identification. Single-copy protein sequences were aligned separately using MAFFT [140] (v7.475), trimmed with trimAl [141] and concatenated into a supermatrix. A maximum-likelihood phylogeny was reconstructed using IQ-TREE 2 [142], with branch support evaluated using 1,000 bootstrap replicates. Divergence times were estimated with MCMCtree [143] using fossil or secondary calibration constraints obtained from TimeTree [144].

#### Gene-family evolution

Gene-family expansion and contraction were inferred using CAFE5 [145] based on orthogroup copy numbers and the time-calibrated ultrametric species tree. Lipid-related gene families were identified according to curated functional annotations and conserved protein domains, and their copy numbers were compared across the seven genomes to characterize lineage-specific changes in lipid-related gene repertoires.

#### Identification and conservation analysis of reproductive-associated genes

Reproductive-associated candidate regions were identified through whole-genome synteny analysis among Dura, HapG, and HapO using SyRI v1.6 [138] and plotsr v1.1.1 [139]. Candidate genes were annotated based on functional information from UniProt/Swiss-Prot and homologous searches performed using DIAMOND v2.1.14 [146]. Protein sequence conservation between HapG and HapO alleles was evaluated using pairwise alignment with BLASTP [114], and sequence identity, coverage, and frameshift mutations were assessed. Reciprocal homology searches and phylogenetic analyses were performed using MAFFT v7 [140] for multiple sequence alignment and IQ-TREE v2 [142] for phylogenetic reconstruction. Structural variation information from SyRI [138] was integrated to evaluate the conservation of candidate loci and the structural context surrounding reproductive-associated genes.

#### Ancestral karyotype reconstruction

Ancestral karyotypes were reconstructed using WGDI [66] based on homologous gene pairs, collinear blocks and synonymous substitution rates. Collinearity and Ks analyses were performed using the -d, -icl, -ks, -bi and -bk modules. The relationships between reconstructed ancestral chromosomes and extant chromosomes were determined and visualized using the -km module.

### Hi-C analysis of TADs and A/B compartments

#### TAD identification

Quality-filtered Hi-C reads were independently mapped to HapG and HapO, and corrected chromatin-contact matrices were generated at 40-kb resolution. TADs were identified using hicFindTADs in HiCExplorer [147]. The TAD separation score was calculated across each chromosome, significant local minima were defined as TAD boundaries, and intervals between adjacent boundaries were designated as TADs.

#### A/B compartment identification

For compartment analysis, balanced Hi-C contact matrices were generated at 150-kb resolution and analyzed using eigs-cis in cooltools [148]. Because eigenvector signs are arbitrary, the first eigenvector was oriented independently for each chromosome according to its correlation with gene density. The more gene-rich state was designated as the active A compartment and assigned positive values, whereas the less gene-rich state was designated as the inactive B compartment and assigned negative values.

#### Structural comparison of introgressed regions

Homologous introgressed intervals in HapG and HapO were paired according to their syntenic coordinates. Paired TADs were classified as conserved or reconfigured based on their boundary correspondence, and A/B compartment states were compared to identify compartment switching. The association between TAD reconfiguration and compartment switching was evaluated using Fisher’s exact test. TAD lengths and gene-expression distributions were compared using two-sided Mann–Whitney U or Wilcoxon signed-rank tests, as appropriate.

### Transcriptome quantification and haplotype-resolved expression analysis

#### Transcriptome sequencing and expression quantification

RNA was extracted from fruit tissues collected across five developmental stages, and strand-specific libraries were sequenced on the Illumina platform. Quality-filtered reads were independently aligned to HapG and HapO using HISAT2 [149] (v2.2.1). Alignments were sorted and indexed with SAMtools [119], and transcript abundance was quantified using StringTie [150] (v3.0.1). Expression was reported as fragments per kilobase of transcript per million mapped reads (FPKM) and transformed as specified for subsequent statistical analyses.

#### Allelic expression bias and developmental trajectories

One-to-one HapG–HapO allelic pairs were integrated with the five-stage expression matrix. At each stage, the allelic-bias index (ABI) was calculated as (E_HapG − E_HapO)/(E_HapG + E_HapO), ranging from −1 (HapO bias) to +1 (HapG bias). Mean ABI, ABI range and the slope of ABI across stages were used to summarize overall bias, dynamic amplitude and directional change, and allelic pairs were assigned to developmental trajectory classes using predefined criteria.

#### Tandem duplication and locus-level expression analysis

Tandemly duplicated genes were identified according to genomic proximity and sequence homology. For loci containing tandem duplicates, copy-level expression values were averaged to estimate representative locus-level expression and to avoid direct inflation by copy number. Expression levels and absolute ABI values were compared between tandemly duplicated and non-tandem loci using two-sided Mann– Whitney U tests to evaluate the effects of tandem duplication on expression output and allelic balance.

## Acknowledgements

We thank the members of the Zhou lab at the Chinese Academy of Tropical Agricultural Sciences and the Agricultural Genomics Institute at Shenzhen, Chinese Academy of Agricultural Sciences for their insightful discussions and technical support throughout this project. We express our sincere gratitude to the Rubber Research Institute of the Chinese Academy of Tropical Agricultural Sciences for providing the plant material used for oil palm genome sequencing and assembly. We also thank Guiqi Bi for experimental guidance and supervision.

## Author contributions

Xiangnian Su, Yanling Peng, and Xuanwen Yang contributed to conceptualization, methodology, investigation, formal analysis, and data curation. Fan Zhang, Qi Xu, Zhiyao Ma, Yang Dong, Lianzhu Zhou, Hui Xue, Xuejing Cao, Zhi Zou, Yiwen Wang, and Yongfeng Zhou contributed to investigation, resources, data generation, and/or data analysis. Xiangnian Su developed the analytical scripts, performed visualization, and drafted the original manuscript. Yanling Peng and Xuanwen Yang contributed to data interpretation and manuscript preparation. Yongfeng Zhou and Xianhai Zeng supervised the study, administered the project, and contributed to funding acquisition. All authors contributed to writing—review and editing, and read and approved the final manuscript.

## Data Availability Statement

The sequencing data and genome-related resources generated in this study have been submitted to the NCBI under BioProject PRJNA1511864, entitled “Haplotype-resolved telomere-to-telomere genome assembly of the interspecific oil palm hybrid ‘Reyou 40’.” Associated accession numbers will be provided upon release. Additional datasets are available from the corresponding author upon reasonable request.

## Supporting Information

Additional supporting information is provided in the accompanying Supporting Information file, including Supplementary Figures S1–S17 and Supplementary Tables S1–S19.

## Supporting Information

**This Supporting Information file includes: (detailed in separate EXCEL file)**

- Supplementary Figure S1 to S17 (This file)
- Supplementary Table S1 to S19 (Separate EXCEL file)

## Table of Contents

### Legends for Supplementary Figures S1-S17

**Figure S1** K-mer-based characterization of the interspecific hybrid genome.

**Figure S2** Copy-number distributions and composition of long terminal repeat retrotransposons in HapO and HapG.

**Figure S3** Sequence composition and similarity between HapG and HapO.

**Figure S4** Representative haplotype-specific presence-absence variation gene clusters and their functional bias.

**Figure S5** Contribution of transposable elements and structural variations to haplotype-specific PAV genes.

**Figure S6** Comparative orthogroup analysis among seven monocot genomes.

**Figure S7** Functional characteristics of expanded gene families in HapG and HapO.

**Figure S8** Synonymous-substitution distributions and gene-duplication modes.

**Figure S9** Genome-wide distribution and size characteristics of structural variations among Dura, HapG, and HapO.

**Figure S10** Conservation of reproductive-related genomic regions among oil palm genomes and date palm.

**Figure S11** Pairwise synteny surrounding the six candidate introgressed regions in HapG.

**Figure S12** Evolutionary characteristics of the six candidate introgressed regions in HapG.

**Figure S13** Phylogenetic relationships and classification of resistance-related gene families across palms.

**Figure S14** Functional enrichment of genes associated with differentiated and PAV regions.

**Figure S15** Chromatin organization and expression characteristics of candidate introgressed regions.

**Figure S16** Functional enrichment of allelic genes with HapG- or HapO-dominant expression.

**Figure S17** Allelic-bias dynamics and functional enrichment of loci with HapG- or HapO-biased expression.

### Supplementary Tables S1-S7 (detailed in separate EXCEL file)

**Table S1.** K-mer-based genome survey statistics for the F1 interspecific hybrid oil palm “Reyou 40”.

|  |  |  |
| --- | --- | --- |
| k = 23 |  |  |
| property | min | max |
| Heterozygosity | 3.67659% | 3.7224% |
| Genome Haploid Length | 1,033,584,075 bp | 1,034,672,382 bp |
| Genome Repeat Length | 352,211,604 bp | 352,582,464 bp |
| Genome Unique Length | 681,372,471 bp | 682,089,919 bp |
| Model Fit | 93.0343% | 96.3299% |
| Read Error Rate | 0.250013% | 0.250013% |
| k = 21 |  |  |
| property | min | max |
| Heterozygosity | 3.61775% | 3.66125% |
| Genome Haploid Length | 974,146,768 bp | 975,203,065 bp |
| Genome Repeat Length | 347,656,722 bp | 348,033,696 bp |
| Genome Unique Length | 626,490,047 bp | 627,169,368 bp |
| Model Fit | 92.8346% | 96.3712% |
| Read Error Rate | 0.244924% | 0.244924% |
| k = 25 |  |  |
| property | min | max |
| Heterozygosity | 3.70823% | 3.75757% |
| Genome Haploid Length | 1,087,023,947 bp | 1,088,145,649 bp |
| Genome Repeat Length | 357,565,818 bp | 357,934,791 bp |
| Genome Unique Length | 729,458,129 bp | 730,210,858 bp |
| Model Fit | 93.2316% | 96.2592% |
| Read Error Rate | 0.251718% | 0.251718% |
| k = 31 |  |  |
| property | min | max |
| Heterozygosity | 3.74507% | 3.81568% |
| Genome Haploid Length | 1,220,825,827 bp | 1,222,016,653 bp |
| Genome Repeat Length | 365,969,342 bp | 366,326,318 bp |
| Genome Unique Length | 854,856,485 bp | 855,690,335 bp |
| Model Fit | 93.8239% | 96.3123% |
| Read Error Rate | 0.247925% | 0.247925% |

**Table S2.**
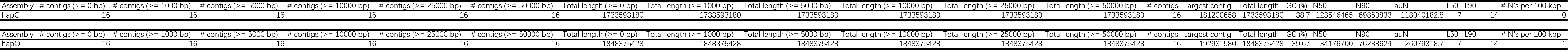
Assembly and BUSCO quality statistics for the haplotype.

| Assembly | # contigs (>= 0 bp) | # contigs (>= 1000 bp) | # contigs (>= 5000 bp) | # contigs (>= 10000 bp) | # contigs (>= 25000 bp) | # contigs (>= 50000 bp) | Total length (>= 0 bp) | Total length (>= 1000 bp) | Total length (>= 5000 bp) | Total length (>= 10000 bp) | Total length (>= 25000 bp) | Total length (>= 50000 bp) | # contigs | Largest contig | Total length | GC (%) | N50 | N90 | auN | L50 | L90 | # N's per 100 kbp |
| --- | --- | --- | --- | --- | --- | --- | --- | --- | --- | --- | --- | --- | --- | --- | --- | --- | --- | --- | --- | --- | --- | --- |
| hapG | 16 | 16 | 16 | 16 | 16 | 16 | 1733593180 | 1733593180 | 1733593180 | 1733593180 | 1733593180 | 1733593180 | 16 | 181200658 | 1733593180 | 38.7 | 123546465 | 69860833 | 118040182.8 | 7 | 14 | 0 |
| Assembly | # contigs (>= 0 bp) | # contigs (>= 1000 bp) | # contigs (>= 5000 bp) | # contigs (>= 10000 bp) | # contigs (>= 25000 bp) | # contigs (>= 50000 bp) | Total length (>= 0 bp) | Total length (>= 1000 bp) | Total length (>= 5000 bp) | Total length (>= 10000 bp) | Total length (>= 25000 bp) | Total length (>= 50000 bp) | # contigs | Largest contig | Total length | GC (%) | N50 | N90 | auN | L50 | L90 | # N's per 100 kbp |
| hapO | 16 | 16 | 16 | 16 | 16 | 16 | 1848375428 | 1848375428 | 1848375428 | 1848375428 | 1848375428 | 1848375428 | 16 | 192931980 | 1848375428 | 39.67 | 134176700 | 76238624 | 126079318.7 | 7 | 14 | 1 |

**Table S3.** Assembly and N50 quality statistics for the haplotype.

| Haplotype | BUSCO category | Abbreviation | Number | Percentage (%) | Notes |
| --- | --- | --- | --- | --- | --- |
| HapG | Complete BUSCOs | C | 2015 | 99.5 | 145 complete BUSCOs contain internal stop codons |
| HapG | Complete and single-copy BUSCOs | S | 1734 | 85.6 | — |
| HapG | Complete and duplicated BUSCOs | D | 281 | 13.9 | — |
| HapG | Fragmented BUSCOs | F | 9 | 0.4 | — |
| HapG | Missing BUSCOs | M | 2 | 0.1 | — |
| HapG | Total BUSCO groups searched | n | 2026 | 100 | — |
| HapO | Complete BUSCOs | C | 2017 | 99.6 | 152 complete BUSCOs contain internal stop codons |
| HapO | Complete and single-copy BUSCOs | S | 1729 | 85.3 | — |
| HapO | Complete and duplicated BUSCOs | D | 288 | 14.2 | — |
| HapO | Fragmented BUSCOs | F | 7 | 0.3 | — |
| HapO | Missing BUSCOs | M | 2 | 0.1 | — |
| HapO | Total BUSCO groups searched | n | 2026 | 100 | — |

**Table S4.**
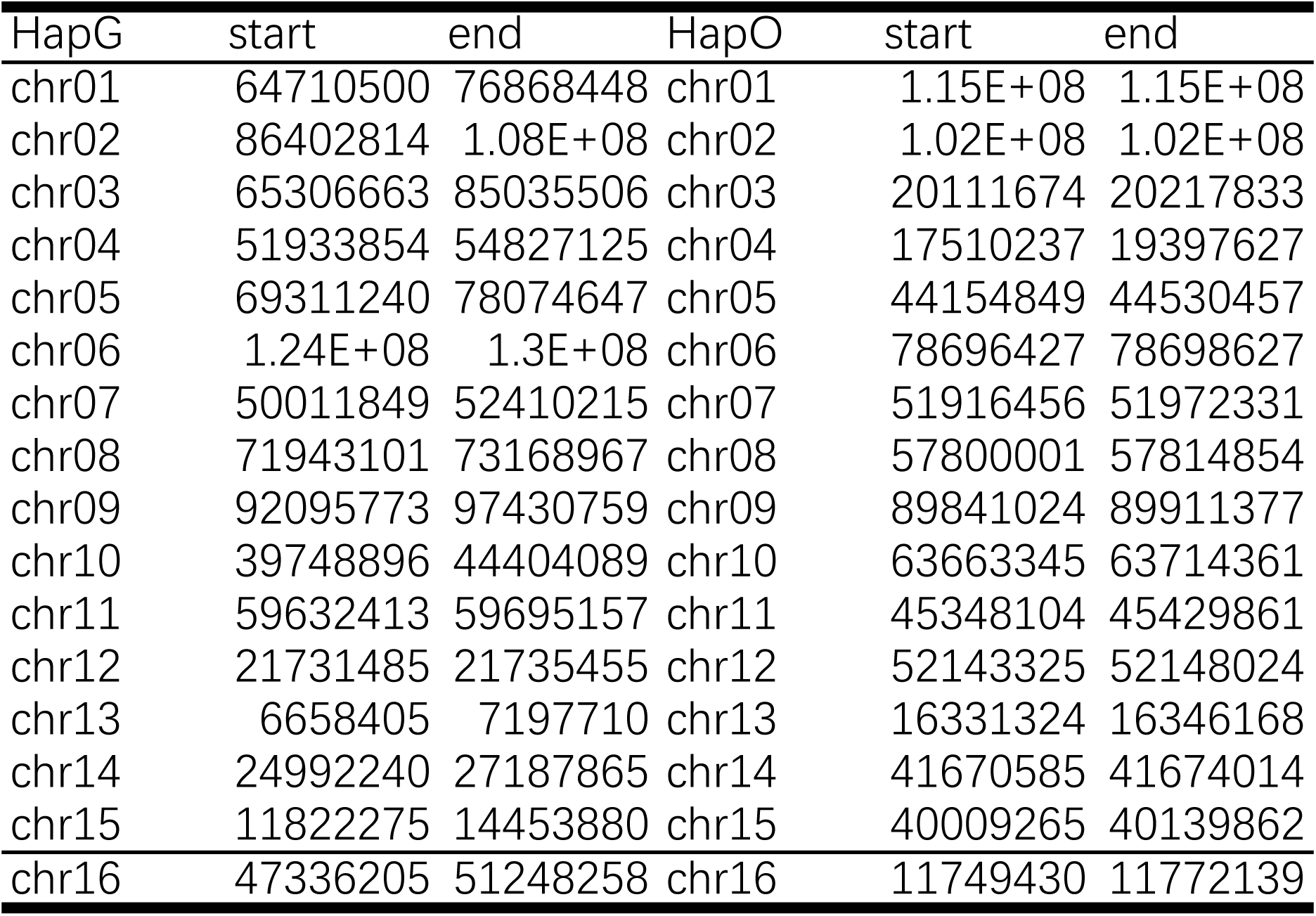
Coordinates and sequence characteristics of CENH3-defined centromeric regions in HapG and HapO.

**Table S5.**
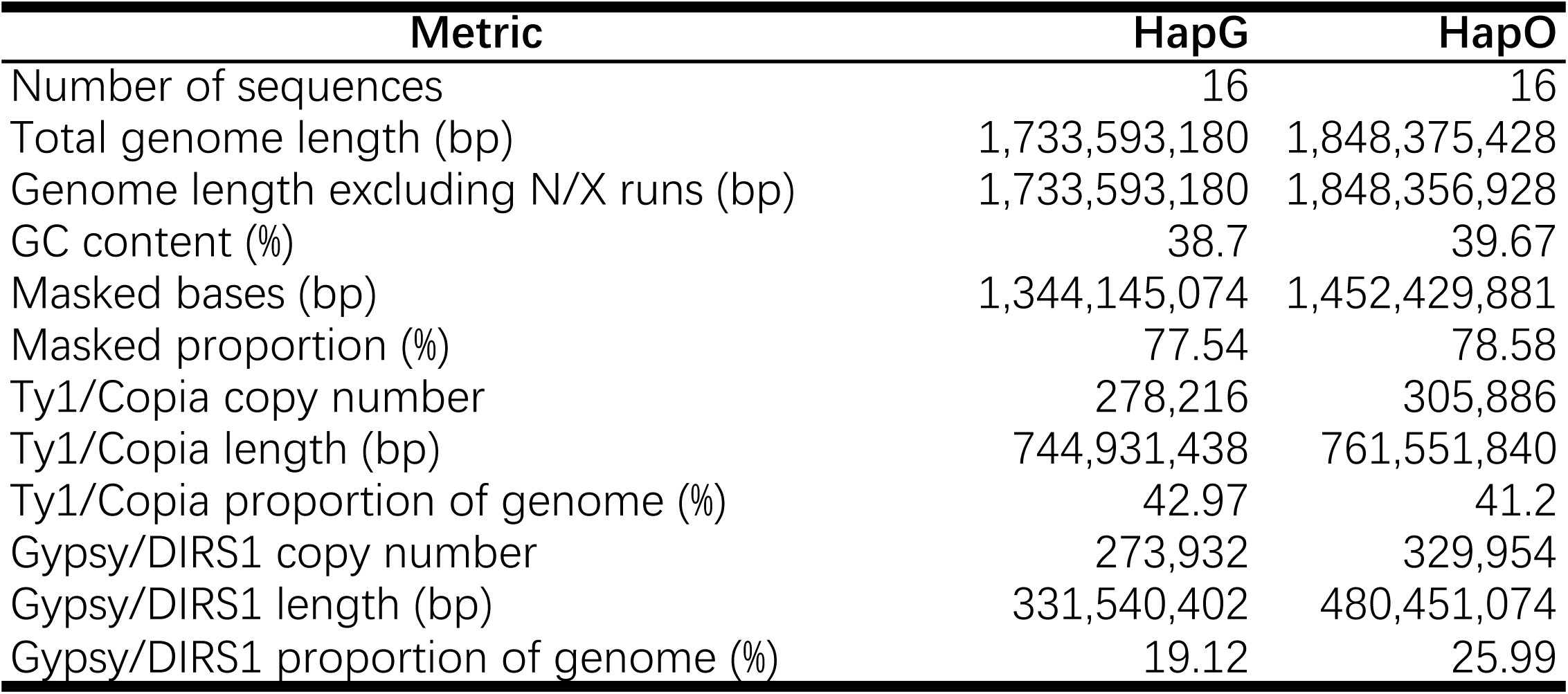
Repeat annotation summary and long terminal repeat retrotransposon composition of HapG and HapO.

| Metric | HapG | HapO |
| --- | --- | --- |
| Number of sequences | 16 | 16 |
| Total genome length (bp) | 1,733,593,180 | 1,848,375,428 |
| Genome length excluding N/X runs (bp) | 1,733,593,180 | 1,848,356,928 |
| GC content (%) | 38.7 | 39.67 |
| Masked bases (bp) | 1,344,145,074 | 1,452,429,881 |
| Masked proportion (%) | 77.54 | 78.58 |
| Ty1/Copia copy number | 278,216 | 305,886 |
| Ty1/Copia length (bp) | 744,931,438 | 761,551,840 |
| Ty1/Copia proportion of genome (%) | 42.97 | 41.2 |
| Gypsy/DIRS1 copy number | 273,932 | 329,954 |
| Gypsy/DIRS1 length (bp) | 331,540,402 | 480,451,074 |
| Gypsy/DIRS1 proportion of genome (%) | 19.12 | 25.99 |

**Table S6.**
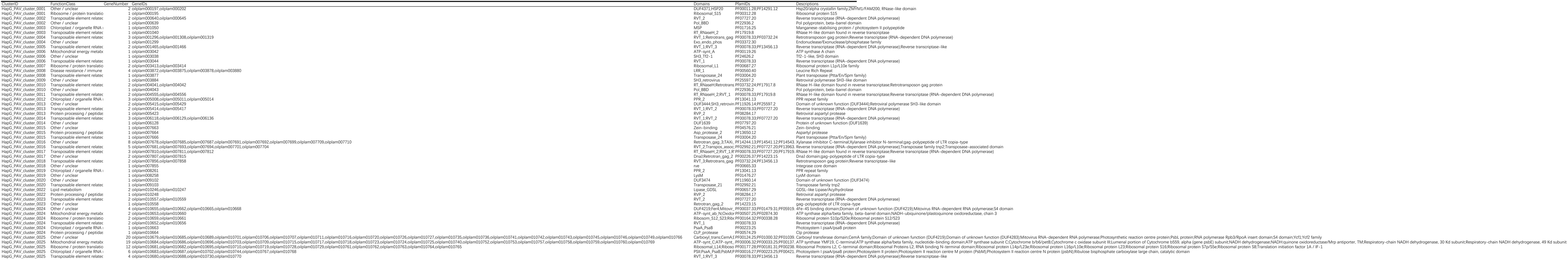
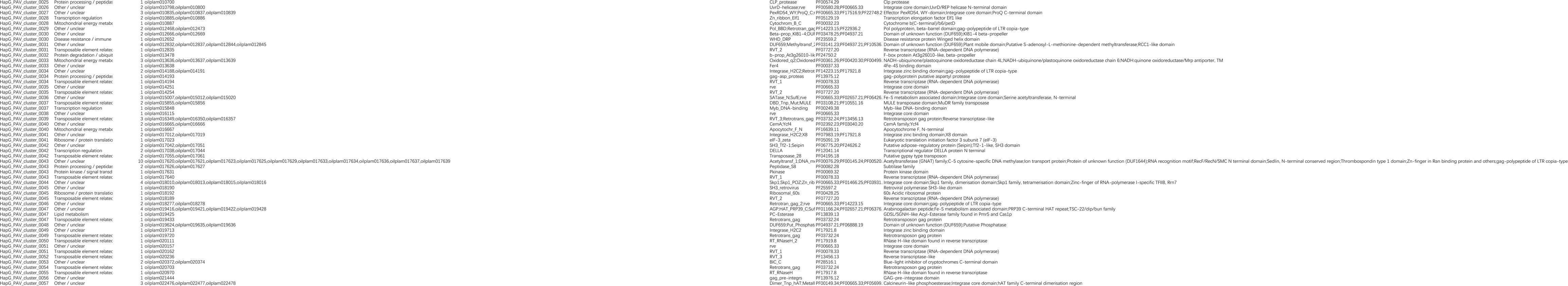
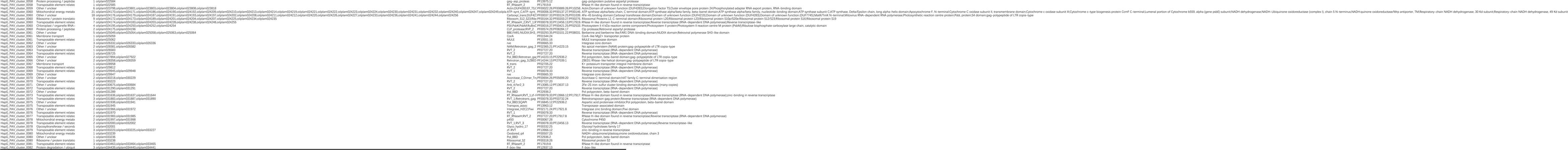
HapG-specific presence-absence variation genes and genomic PAV clusters.

**Table S7.**
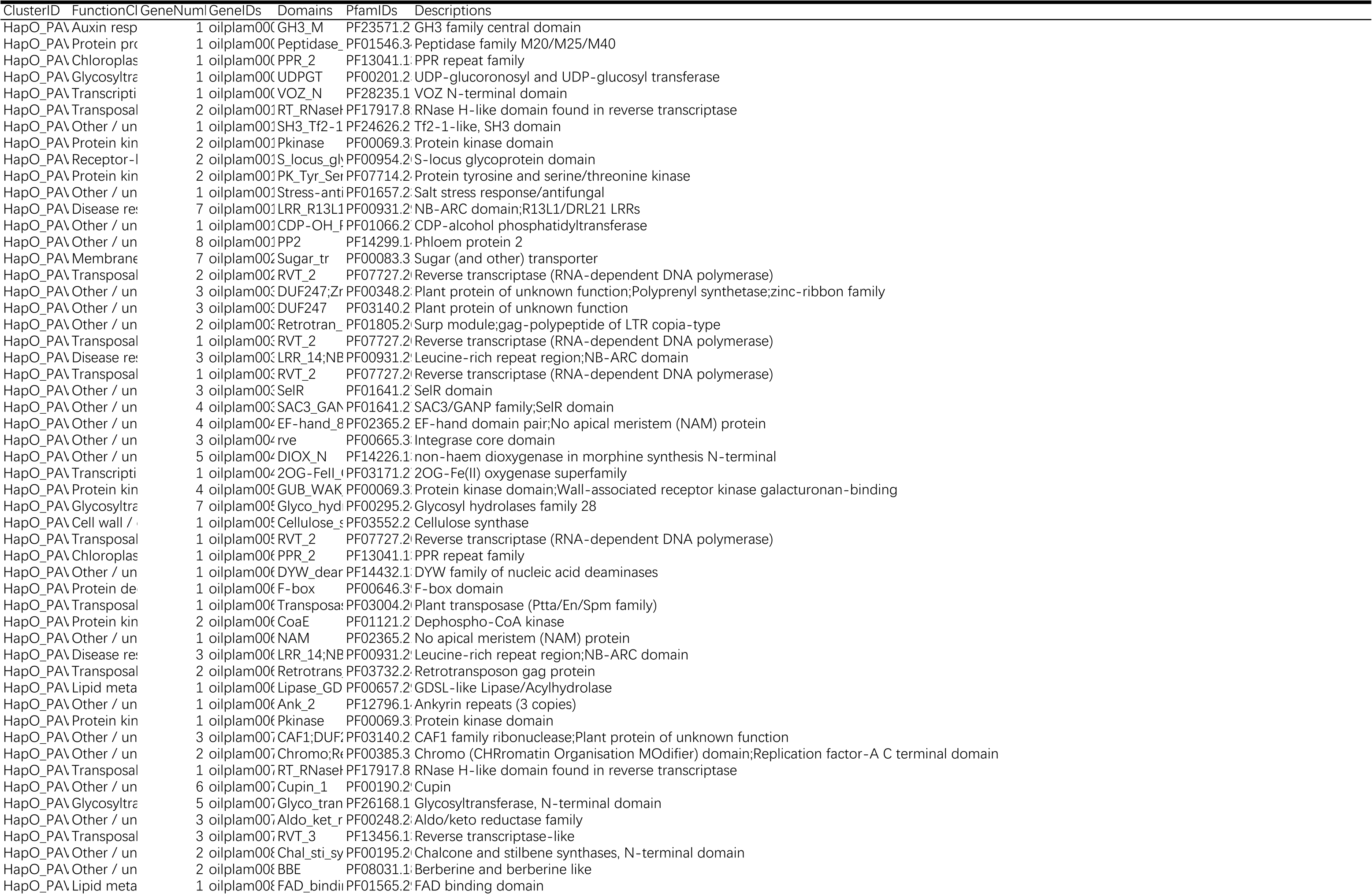
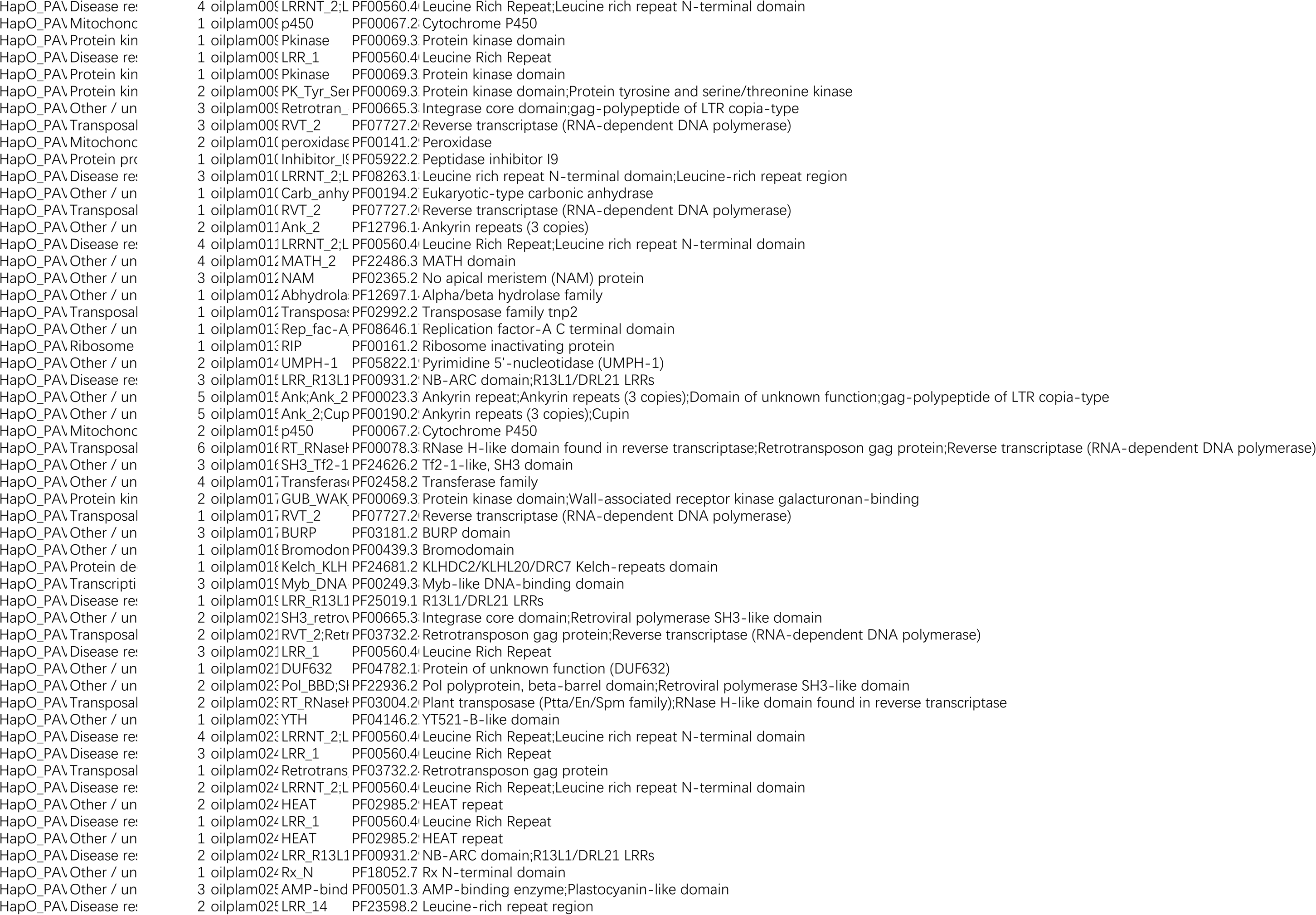
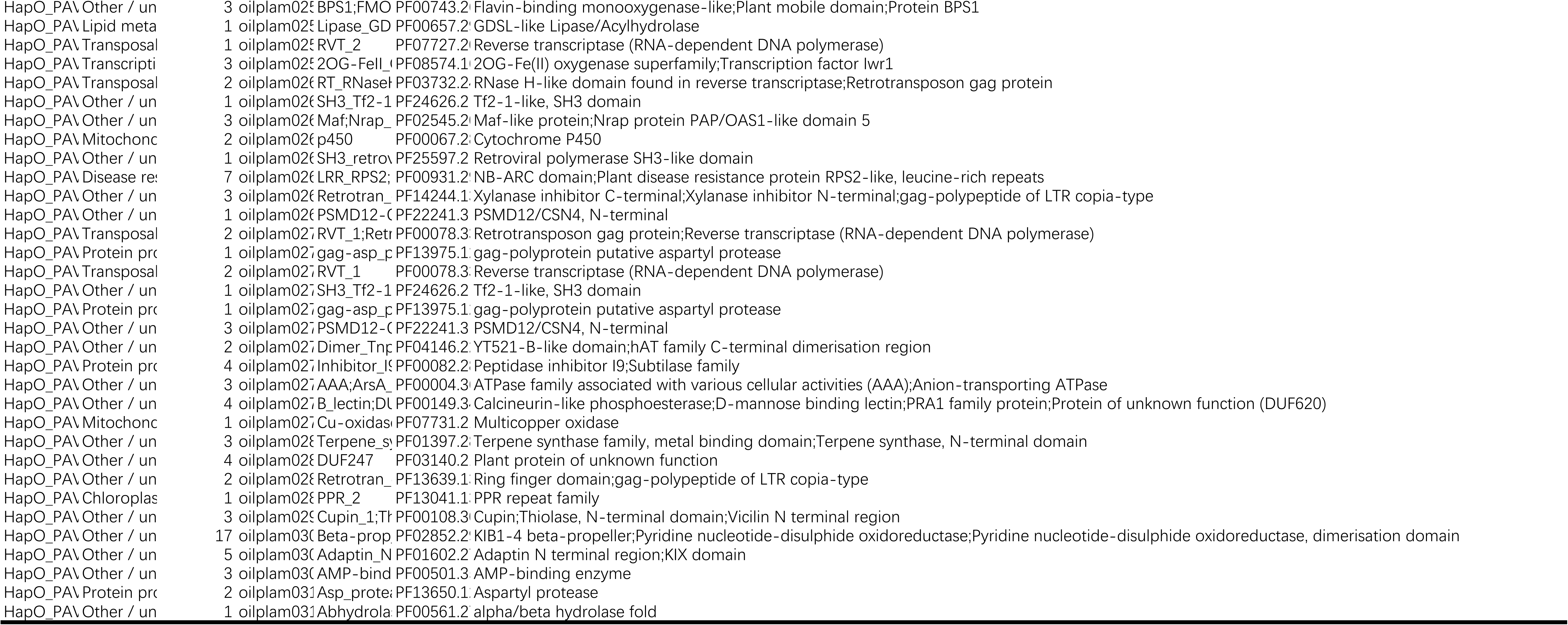
HapO-specific presence-absence variation genes and genomic PAV clusters.

**Table S8.** Chromosome-scale synteny and collinearity statistics among HapG, HapO, and representative palm genomes.

| Species_pair | File | Syntenic_blocks |
| --- | --- | --- |
| *Cocos nucifera* – *Areca catechu* | Cocos nucifera – Areca catechu.anchors.simple | 321 |
| *Elaeis oleifera* – *Cocos nucifera* | Elaeis oleifera – Cocos nucifera.anchors.simple | 329 |
| *Elaeis guineensis* – *Cocos nucifera* | Elaeis guineensis – Cocos nucifera.anchors.simple | 343 |
| *Areca catechu* – *Elaeis oleifera* | Areca catechu – Elaeis oleifera.anchors.simple | 354 |
| *Phoenix dactylifera* – *Cocos nucifera* | Phoenix dactylifera – Cocos nucifera.anchors.simple | 359 |
| *Cocos nucifera* – *Asparagus officinalis* | Cocos nucifera – Asparagus officinalis.anchors.simple | 405 |
| *Phoenix dactylifera* – *Elaeis oleifera* | Phoenix dactylifera – Elaeis oleifera.anchors.simple | 428 |
| *Phoenix dactylifera* – *Elaeis guineensis* | Phoenix dactylifera – Elaeis guineensis.anchors.simple | 433 |
| *Asparagus officinalis* – *Phoenix dactylifera* | Asparagus officinalis – Areca catechu.anchors.simple | 438 |
| *Asparagus officinalis* – *Areca catechu* | Asparagus officinalis – Areca catechu.anchors.simple | 469 |
| *Elaeis oleifera* – *Elaeis guineensis* | Elaeis oleifera – Elaeis guineensis.anchors.simple | 1172 |

**Table S9.**
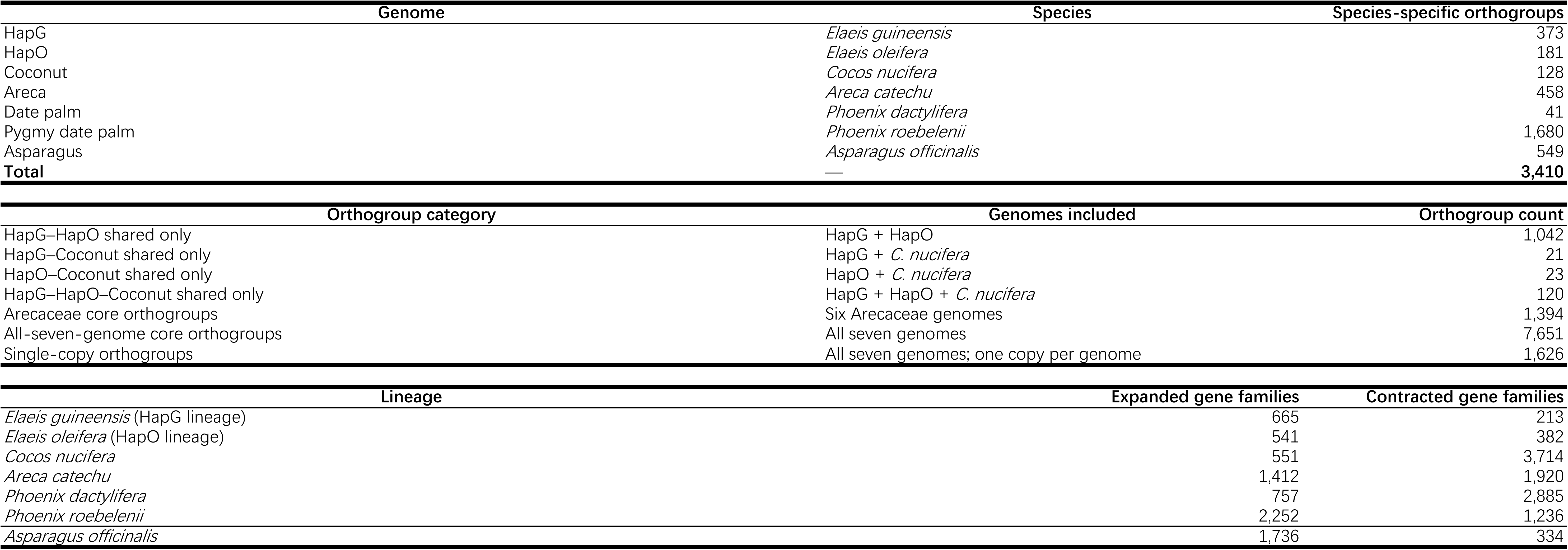
Orthogroup assignments and shared and species-specific gene-family statistics across the seven analyzed genomes.

**Table S10.**
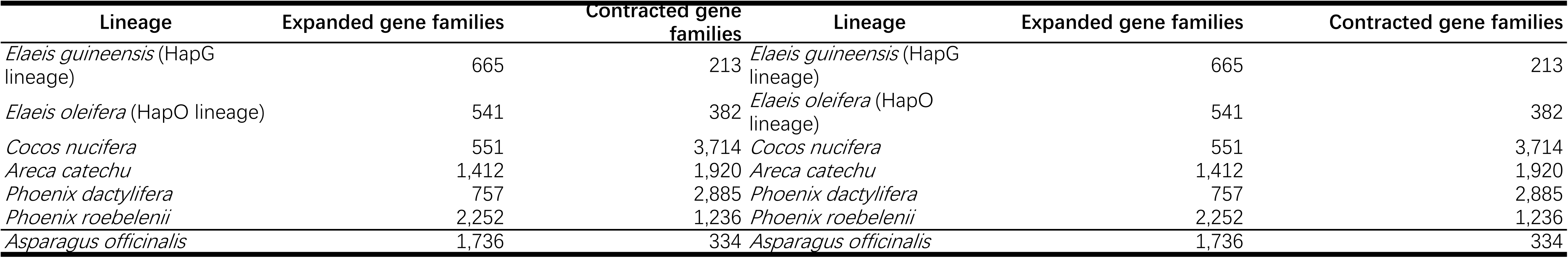
Gene-family expansion and contraction across the time-calibrated palm phylogeny.

**Table S11.**
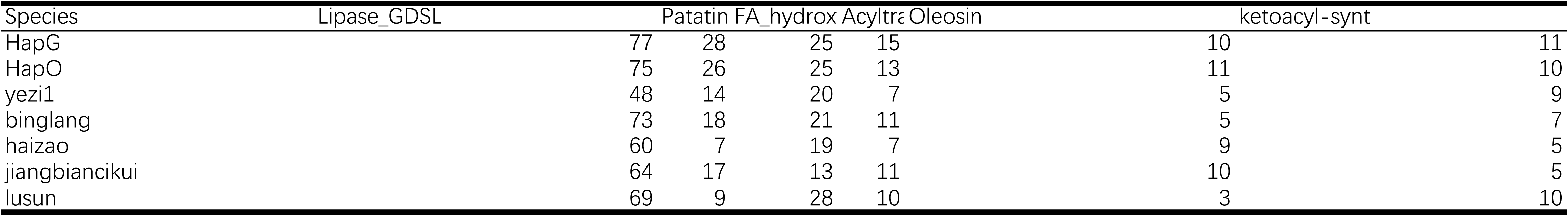
Copy numbers and functional annotations of 17 lipid-related gene families across the analyzed species.

**Table S12.**
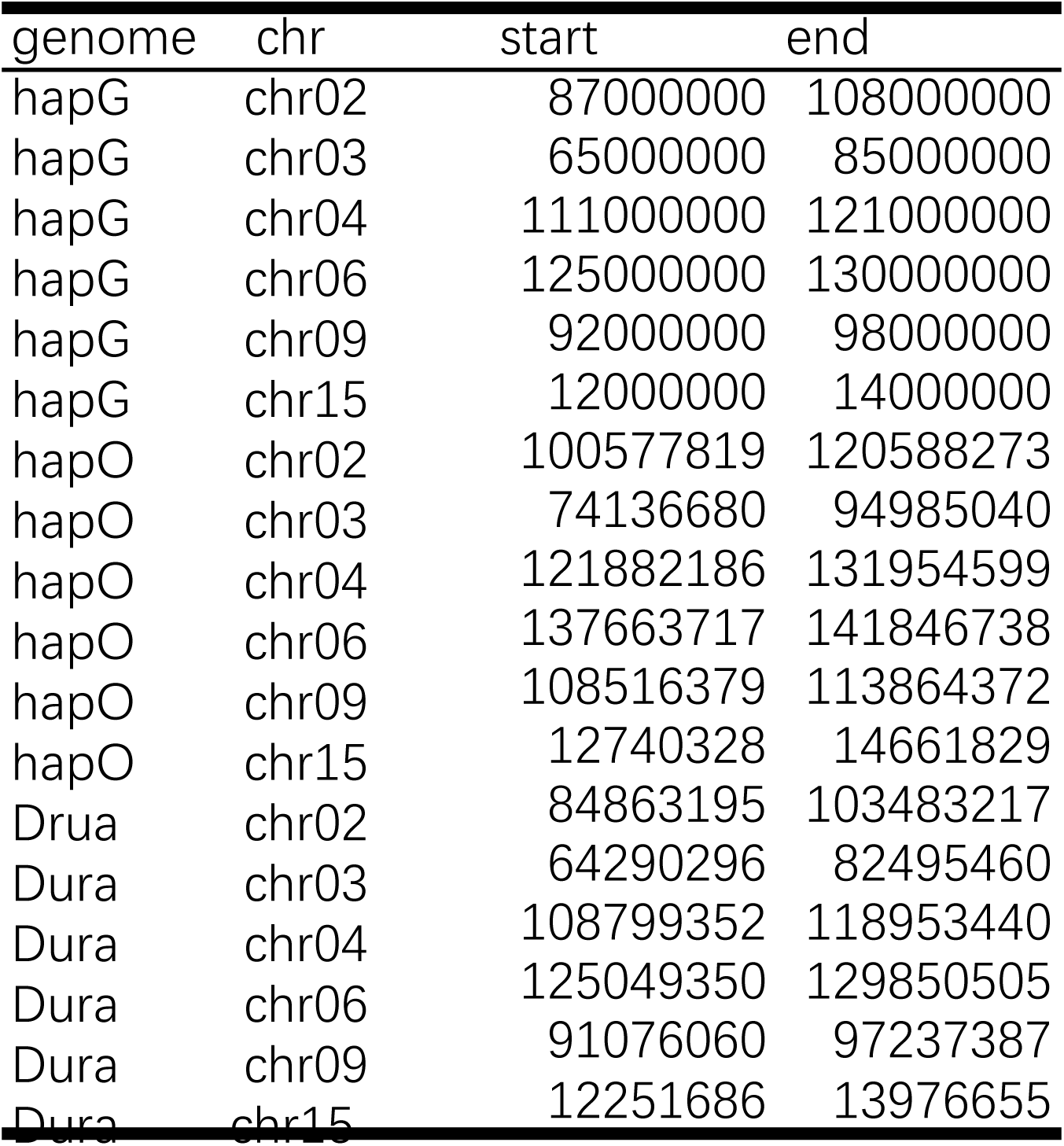
Coordinates, sizes, and summary characteristics of the six candidate ancient introgressed regions in HapG.

| genome | chr | start | end |
| --- | --- | --- | --- |
| hapG | chr02 | 87000000 | 108000000 |
| hapG | chr03 | 65000000 | 85000000 |
| hapG | chr04 | 111000000 | 121000000 |
| hapG | chr06 | 125000000 | 130000000 |
| hapG | chr09 | 92000000 | 98000000 |
| hapG | chr15 | 12000000 | 14000000 |
| hapO | chr02 | 100577819 | 120588273 |
| hapO | chr03 | 74136680 | 94985040 |
| hapO | chr04 | 121882186 | 131954599 |
| hapO | chr06 | 137663717 | 141846738 |
| hapO | chr09 | 108516379 | 113864372 |
| hapO | chr15 | 12740328 | 14661829 |
| Drua | chr02 | 84863195 | 103483217 |
| Dura | chr03 | 64290296 | 82495460 |
| Dura | chr04 | 108799352 | 118953440 |
| Dura | chr06 | 125049350 | 129850505 |
| Dura | chr09 | 91076060 | 97237387 |
| Dura | chr15 | 12251686 | 13976655 |

**Table S13.**
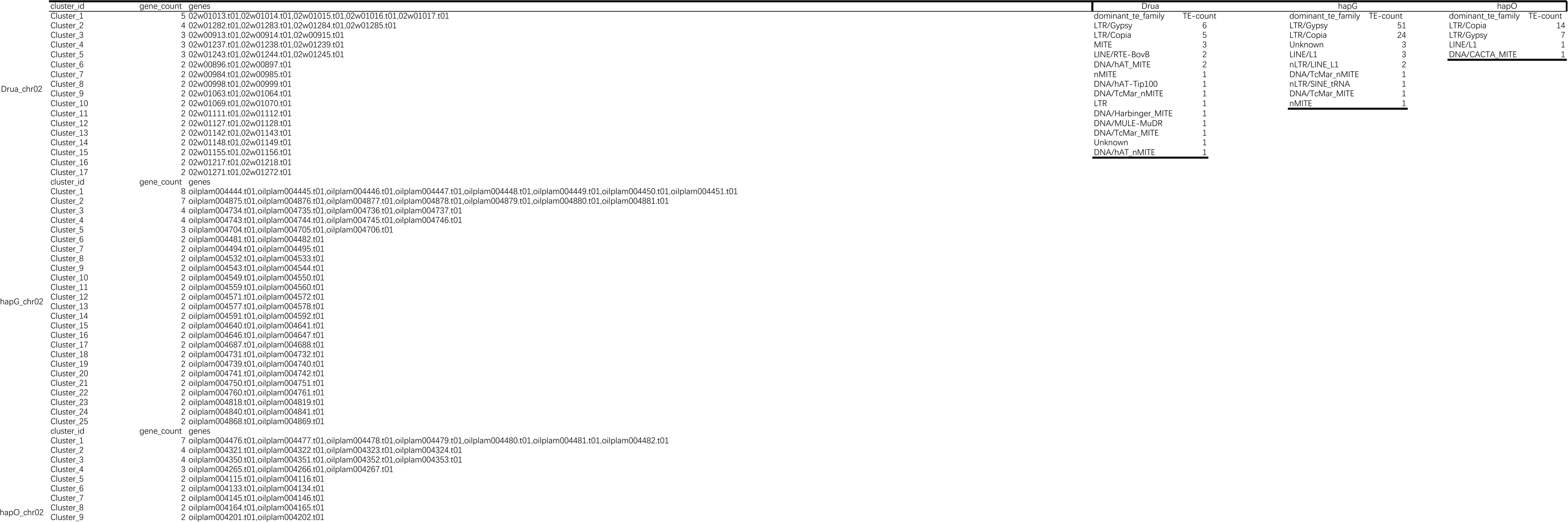
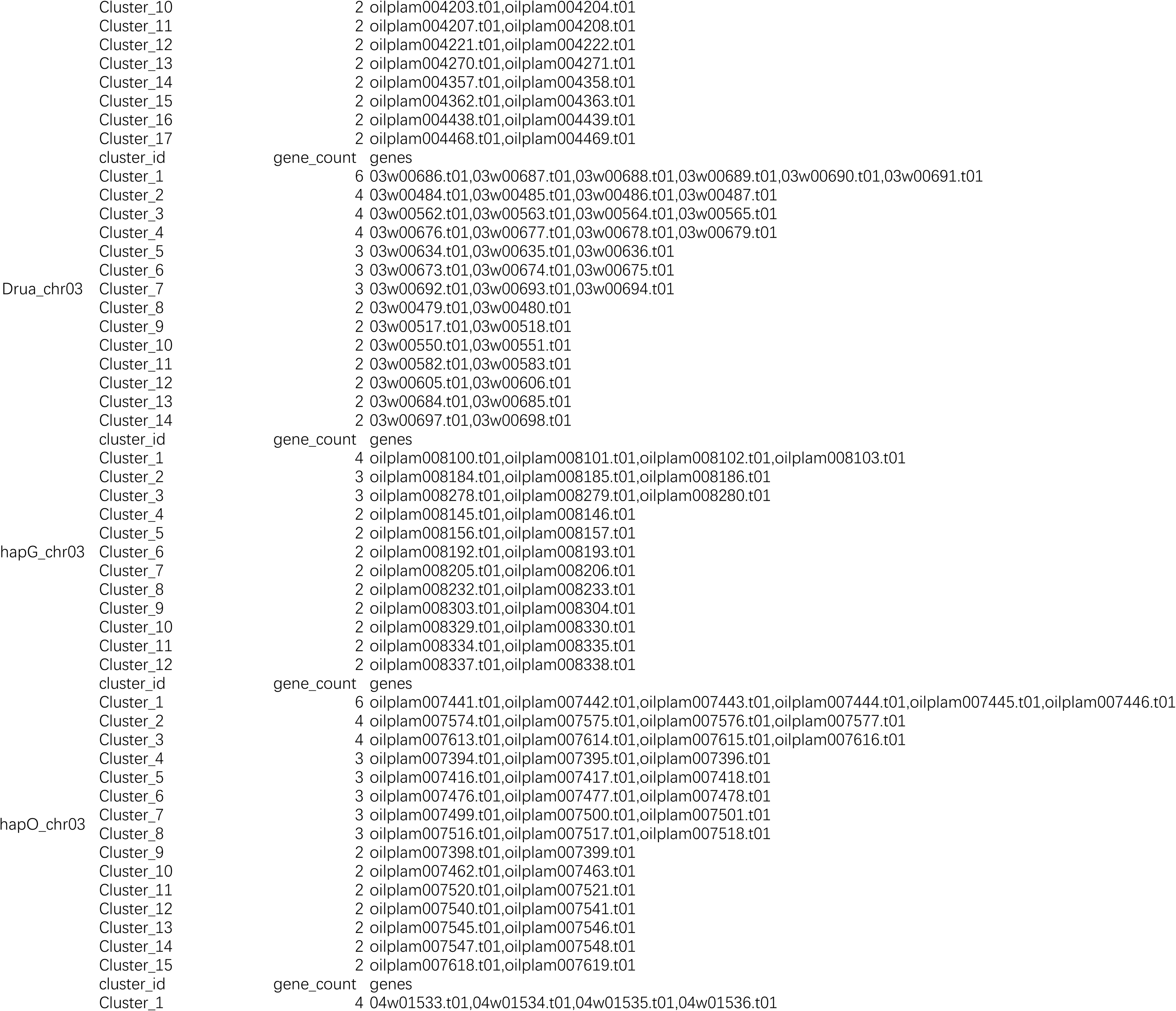
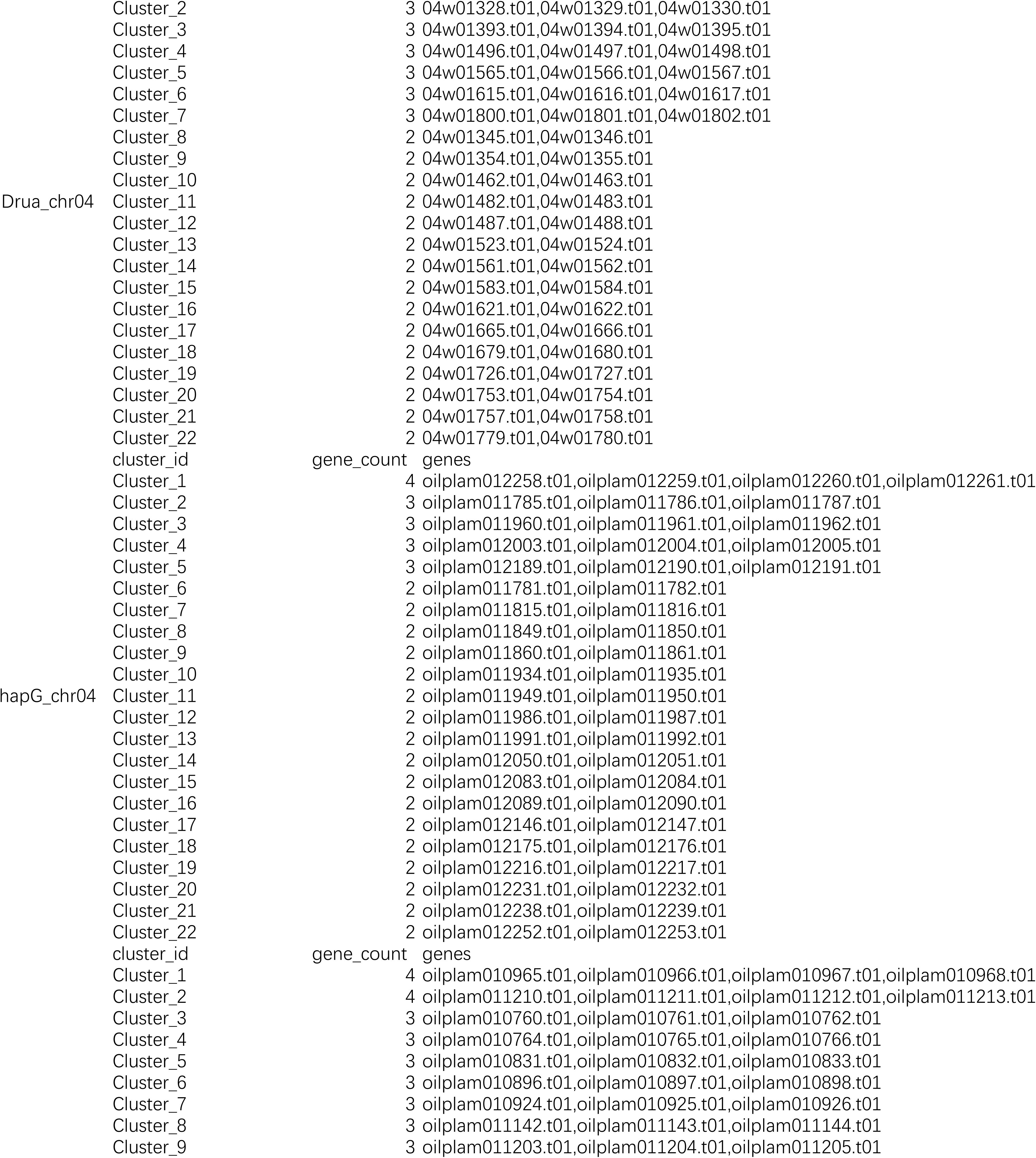
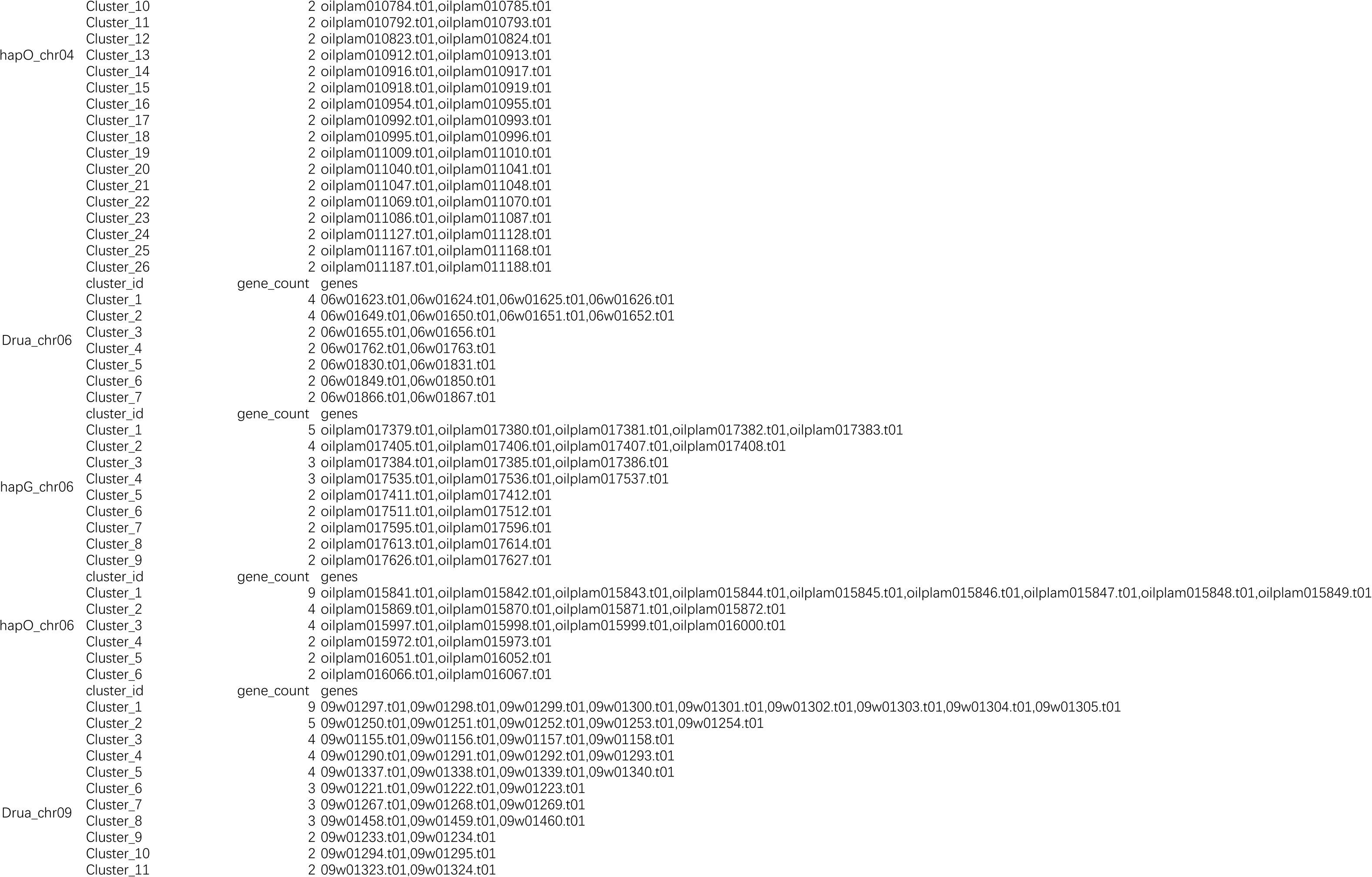
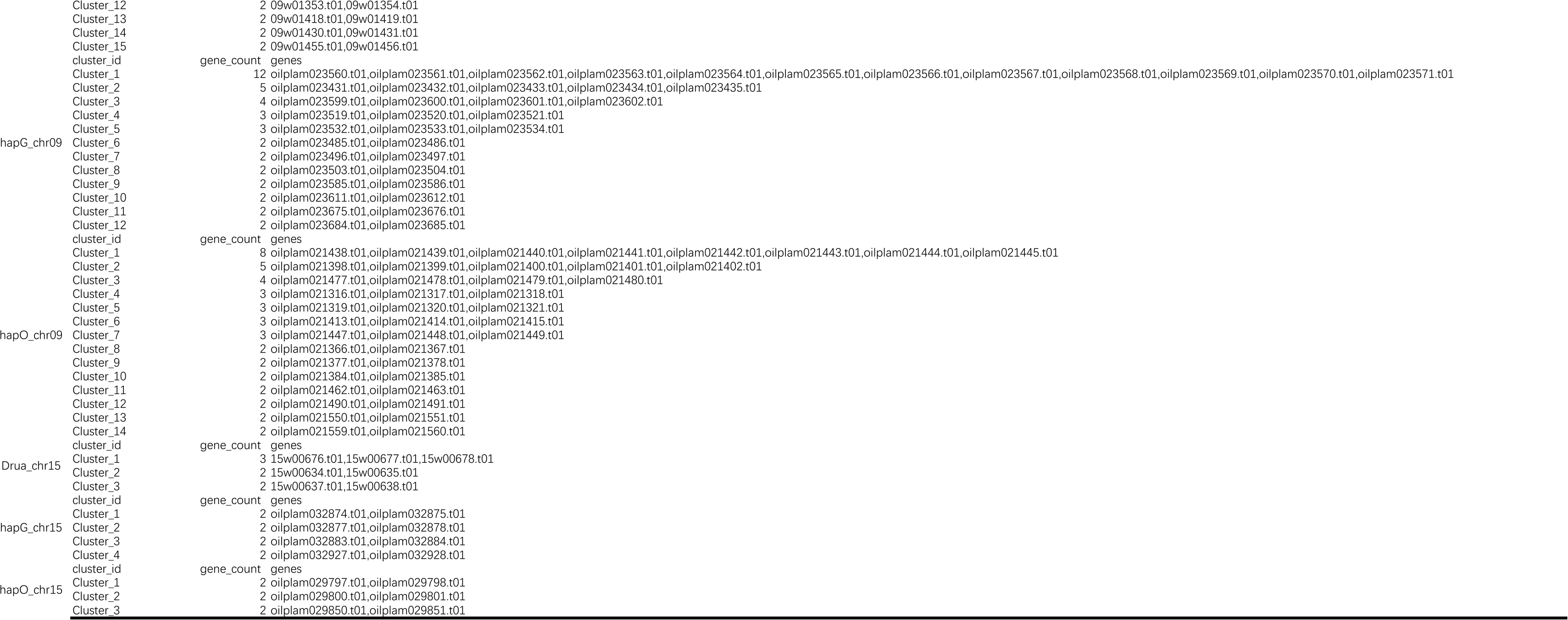
Comparative gene content, tandem duplication, and repeat composition of the six HapG introgressed regions and homologous HapO and Dura intervals.

**Table S14.**
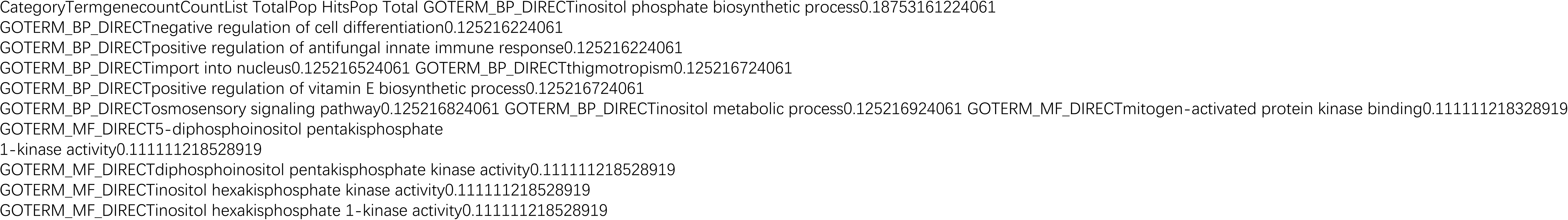
Functional annotations and enrichment of genes associated with highly differentiated segments and presence-absence variation in the candidate introgressed regions.

**Table S15.**
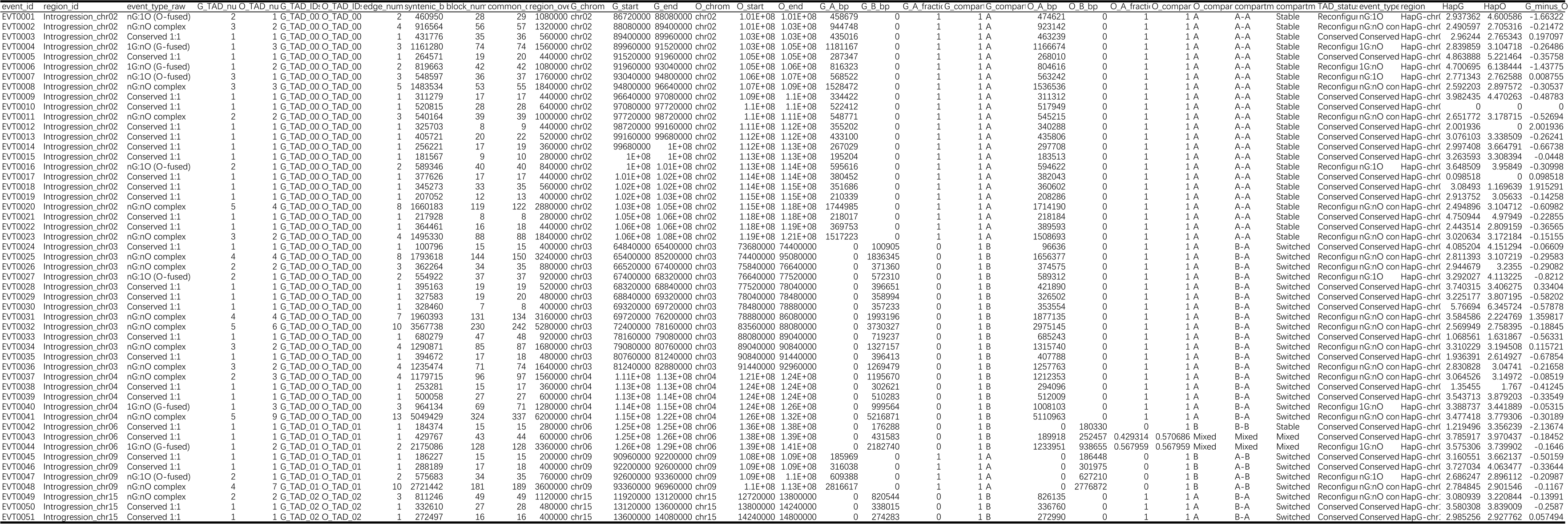
Paired TAD configurations, A/B compartment states, and reconfiguration characteristics of the six candidate introgressed regions.

**Table S16.**
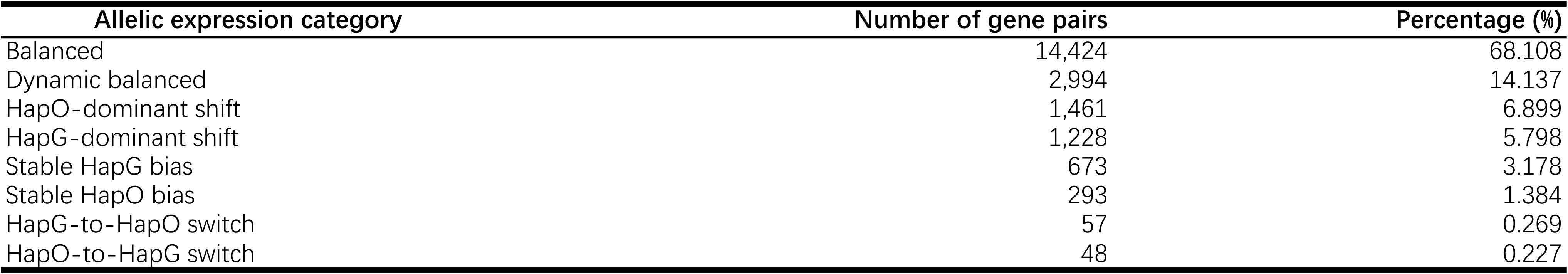
Classification and functional characteristics of balanced and haplotype-biased allelic gene pairs.

**Table S17.**
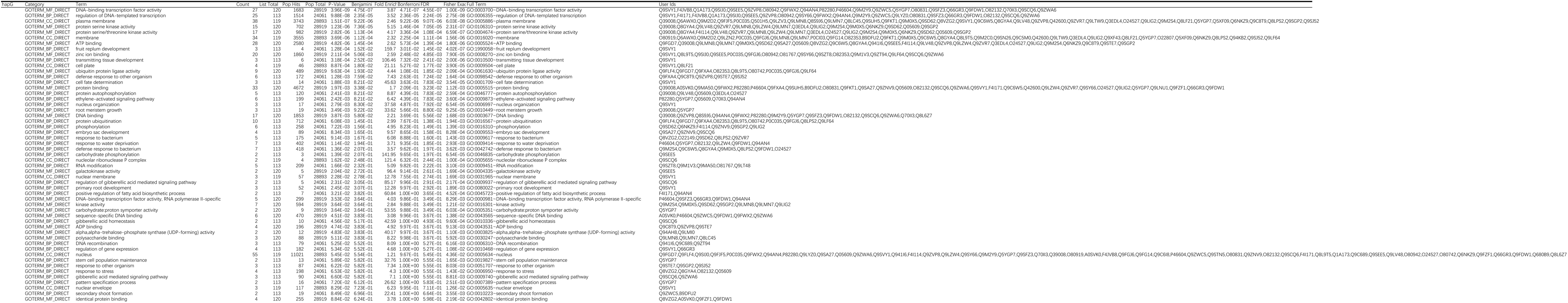
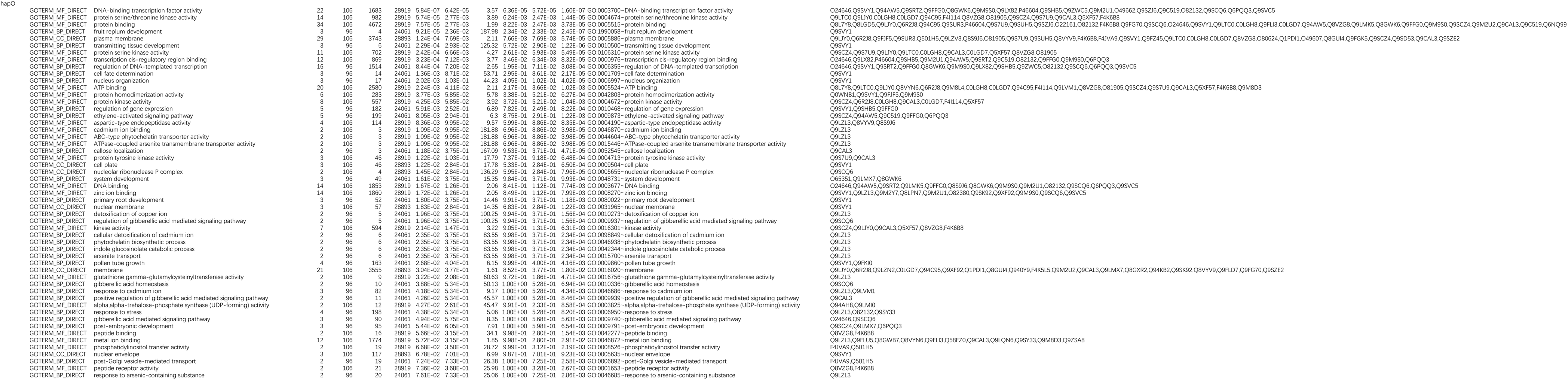
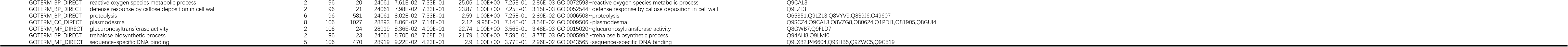
Developmental allelic-bias trajectory classes and functional enrichment across five fruit developmental stages.

**Table S18.**
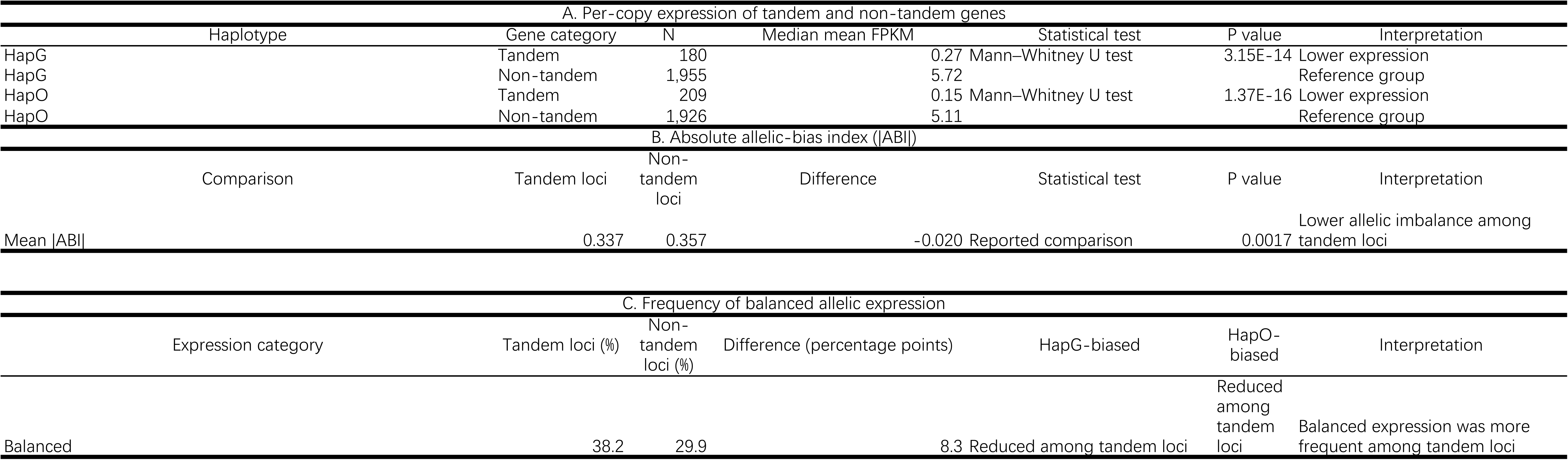
Per-copy expression comparison between tandemly duplicated and non-tandem genes in HapG and HapO.

**Table S19.**
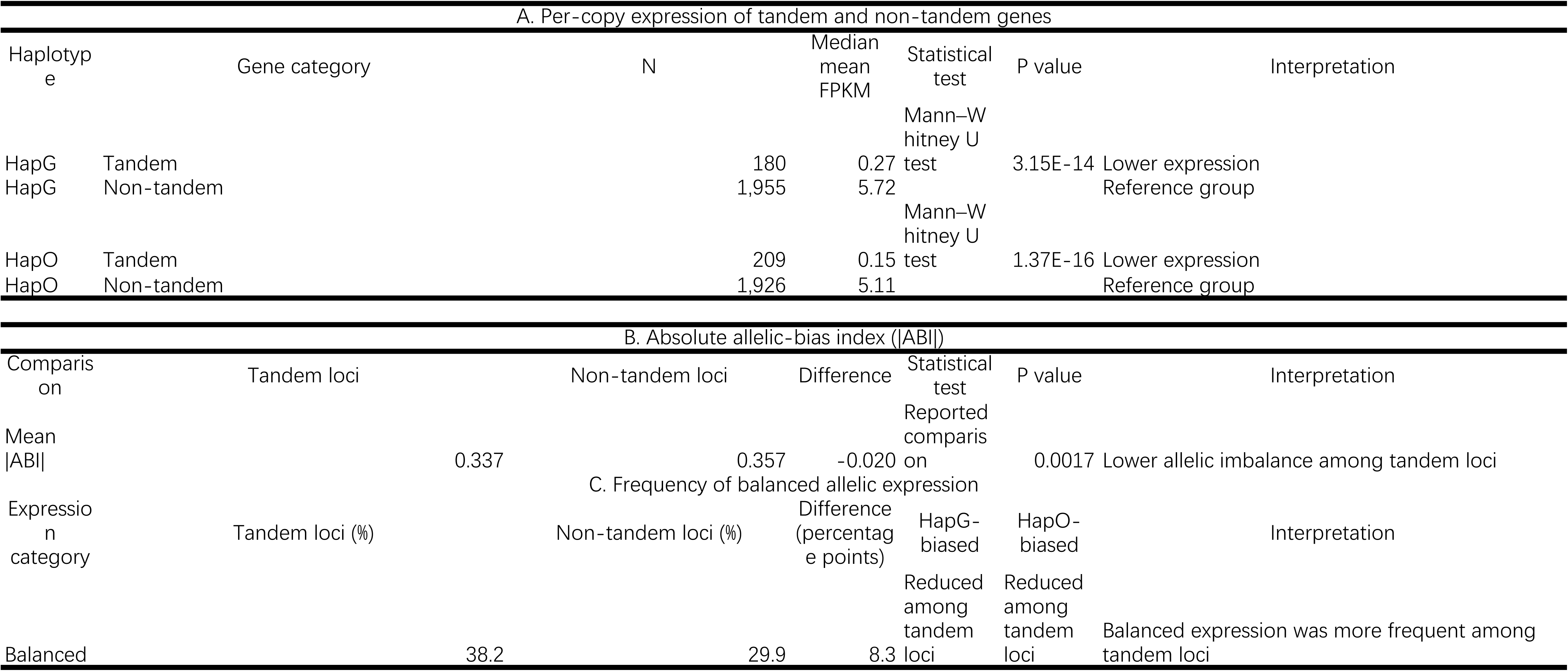
Distribution of HapG-biased, balanced, and HapO-biased loci between tandem and non-tandem groups.

**Abbreviations:** ABI, allelic-bias index; CDS, coding sequence; GO, Gene Ontology; LTR-RT, long terminal repeat retrotransposon; PAV, presence-absence variation; SV, structural variation; TAD, topologically associating domain; TE, transposable element; WGD, whole-genome duplication.

---

**Figure S1.**
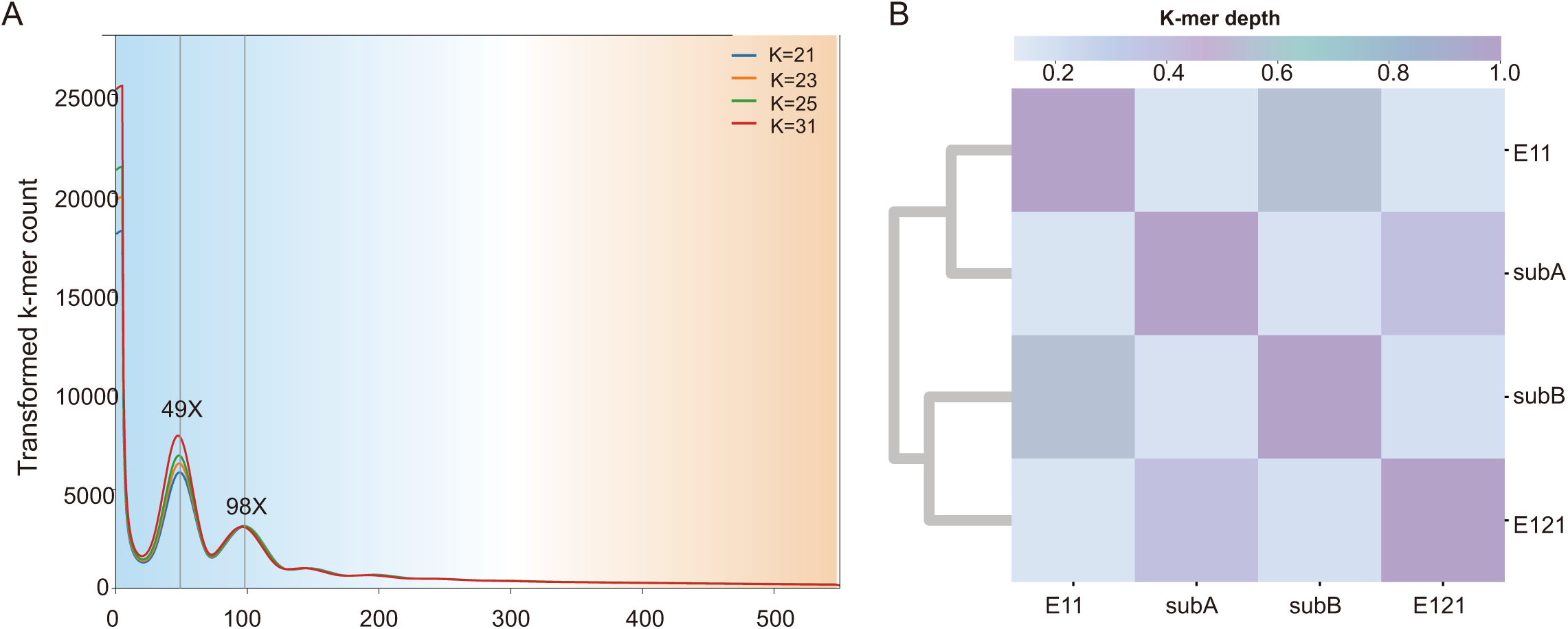
K-mer-based characterization of the interspecific hybrid genome. (A) Transformed k-mer count distributions calculated using k = 21, 23, 25, and 31, showing major depth peaks at approximately 49× and 98×. (B) Heatmap with hierarchical clustering showing k-mer-depth relationships among E11, subA, subB, and E121. The color scale indicates normalized k-mer-depth values.

**Figure S2.**
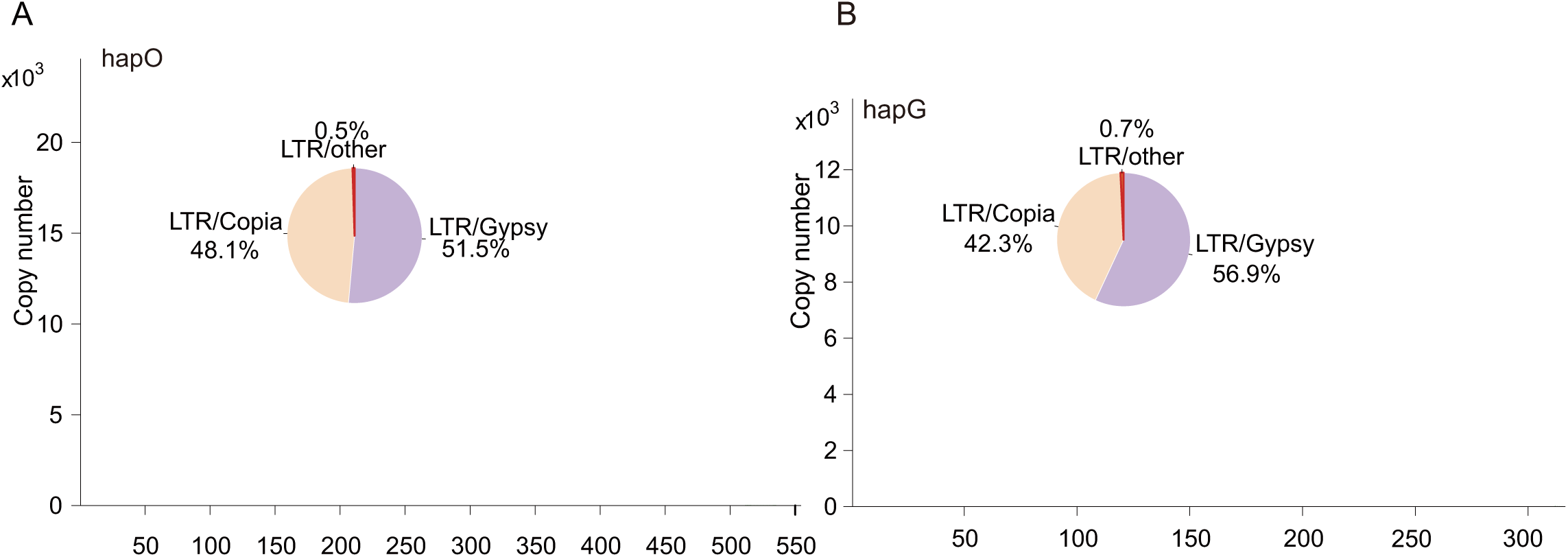
Copy-number distributions and composition of long terminal repeat retrotransposons in HapO and HapG. (A) Long terminal repeat retrotransposon (LTR-RT) copy-number distribution in HapO. The inset shows the relative proportions of LTR/Gypsy (51.5%), LTR/Copia (48.1%), and other LTR elements (0.5%). (B) LTR-RT copy-number distribution in HapG. The inset shows the relative proportions of LTR/Gypsy (56.9%), LTR/Copia (42.3%), and other LTR elements (0.7%).

**Figure S3.**
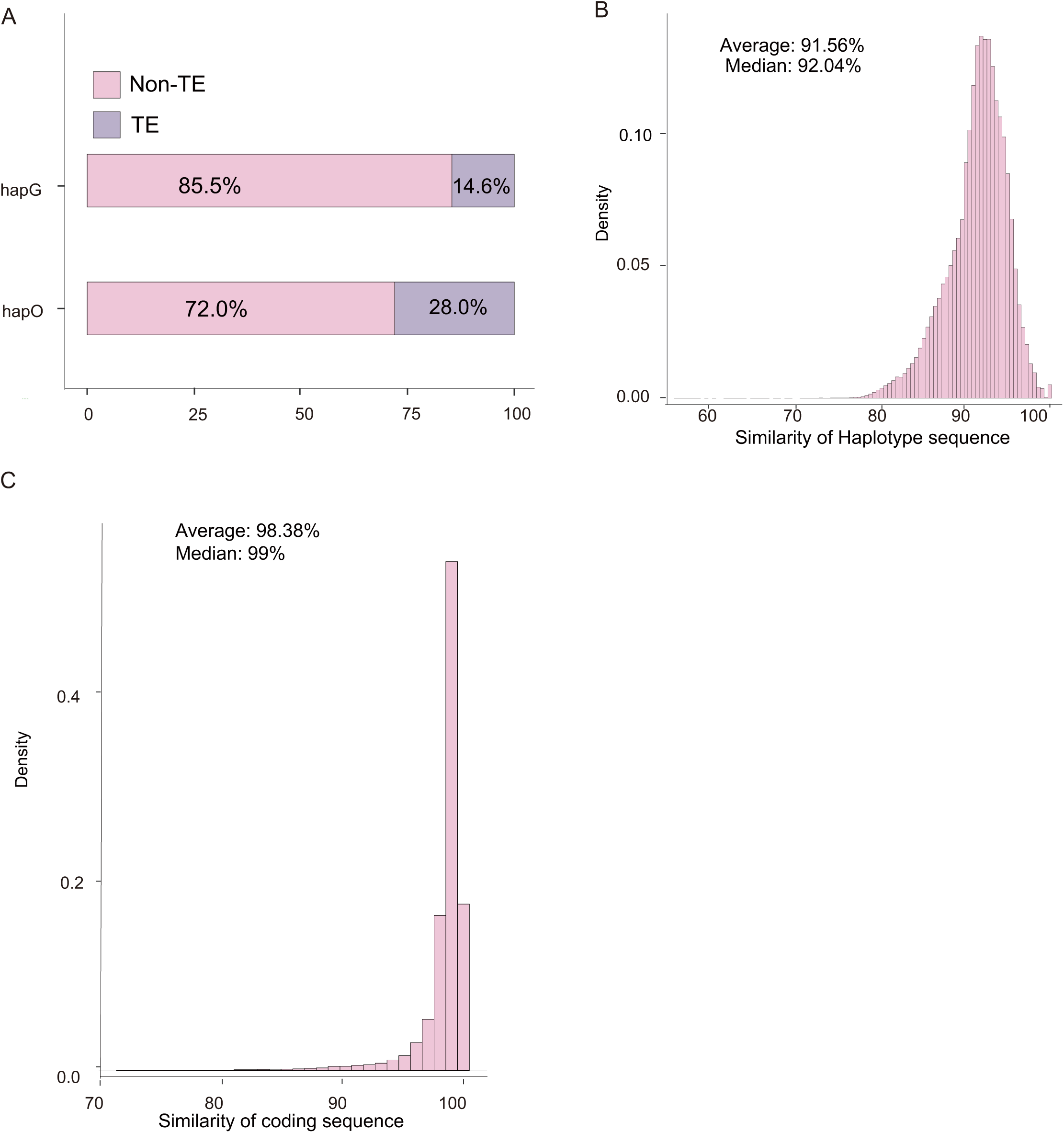
Sequence composition and similarity between HapG and HapO. (A) Relative proportions of transposable-element (TE) and non-TE sequence in the analyzed HapG and HapO sequence sets. (B) Density distribution of HapG-HapO sequence similarity, with an average of 91.56% and a median of 92.04%. (C) Density distribution of coding-sequence similarity between orthologous coding sequences, with an average of 98.38% and a median of 99%.

**Figure S4.**
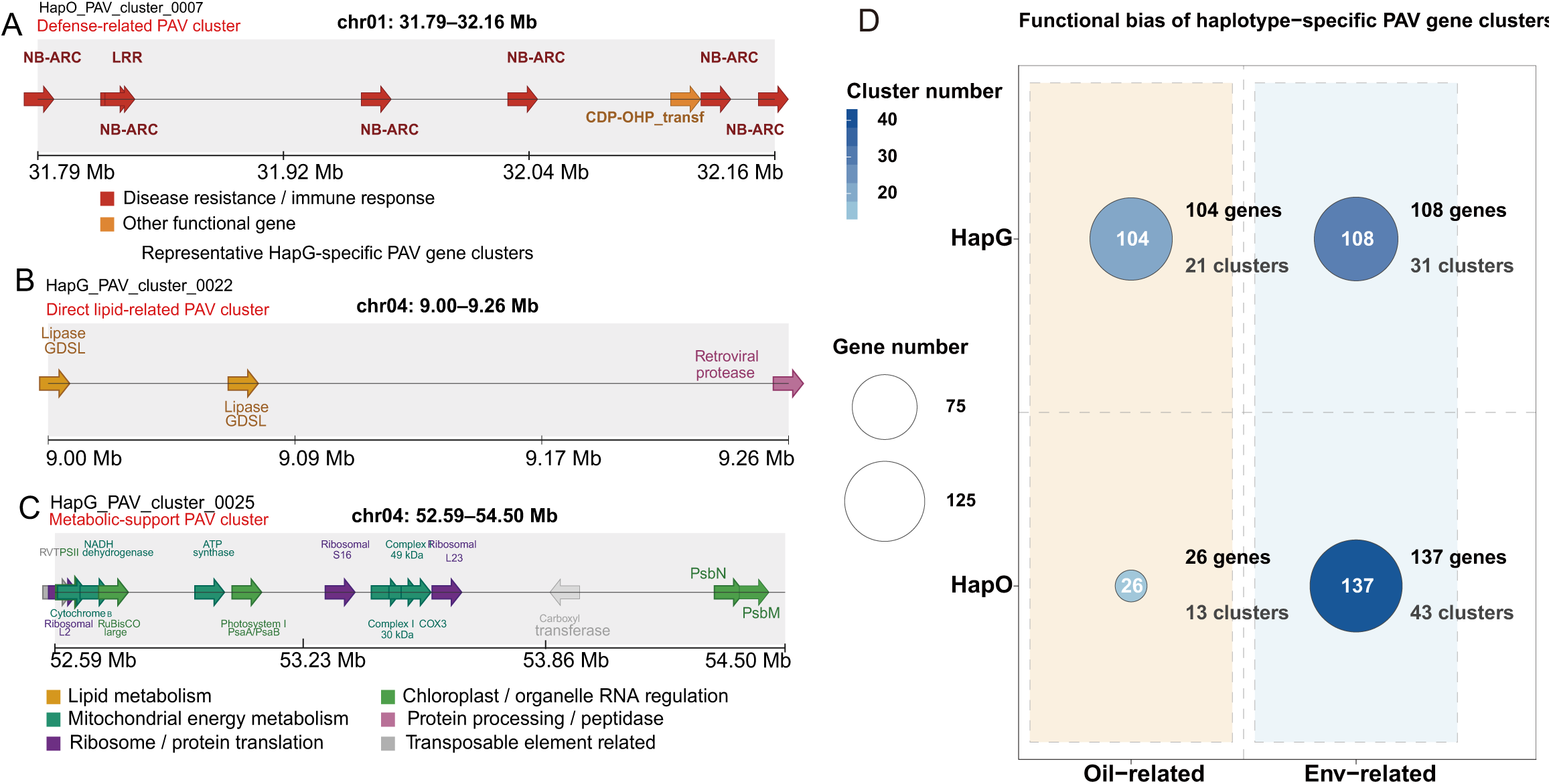
Representative haplotype-specific presence-absence variation gene clusters and their functional bias. (A) HapO_PAV_cluster_0007, a defense-related presence-absence variation (PAV) cluster on chromosome 1 (31.79-32.16 Mb), containing multiple NB-ARC- and LRR-associated genes together with other functional genes. (B) HapG_PAV_cluster_0022, a lipid-related PAV cluster on chromosome 4 (9.00-9.26 Mb), containing GDSL lipase genes and a retroviral-protease-associated gene. (C) HapG_PAV_cluster_0025, a metabolic-support PAV cluster on chromosome 4 (52.59-54.50 Mb), containing genes associated with lipid metabolism, mitochondrial energy metabolism, chloroplast/organelle RNA regulation, ribosome/protein translation, protein processing/peptidase activity, and transposable-element-related functions. (D) Functional bias of haplotype-specific PAV gene clusters toward oil-related and environment-related functions. Bubble size represents gene number and shading represents cluster number. HapG contains 104 oil-related genes in 21 clusters and 108 environment-related genes in 31 clusters; HapO contains 26 oil-related genes in 13 clusters and 137 environment-related genes in 43 clusters.

**Figure S5.**
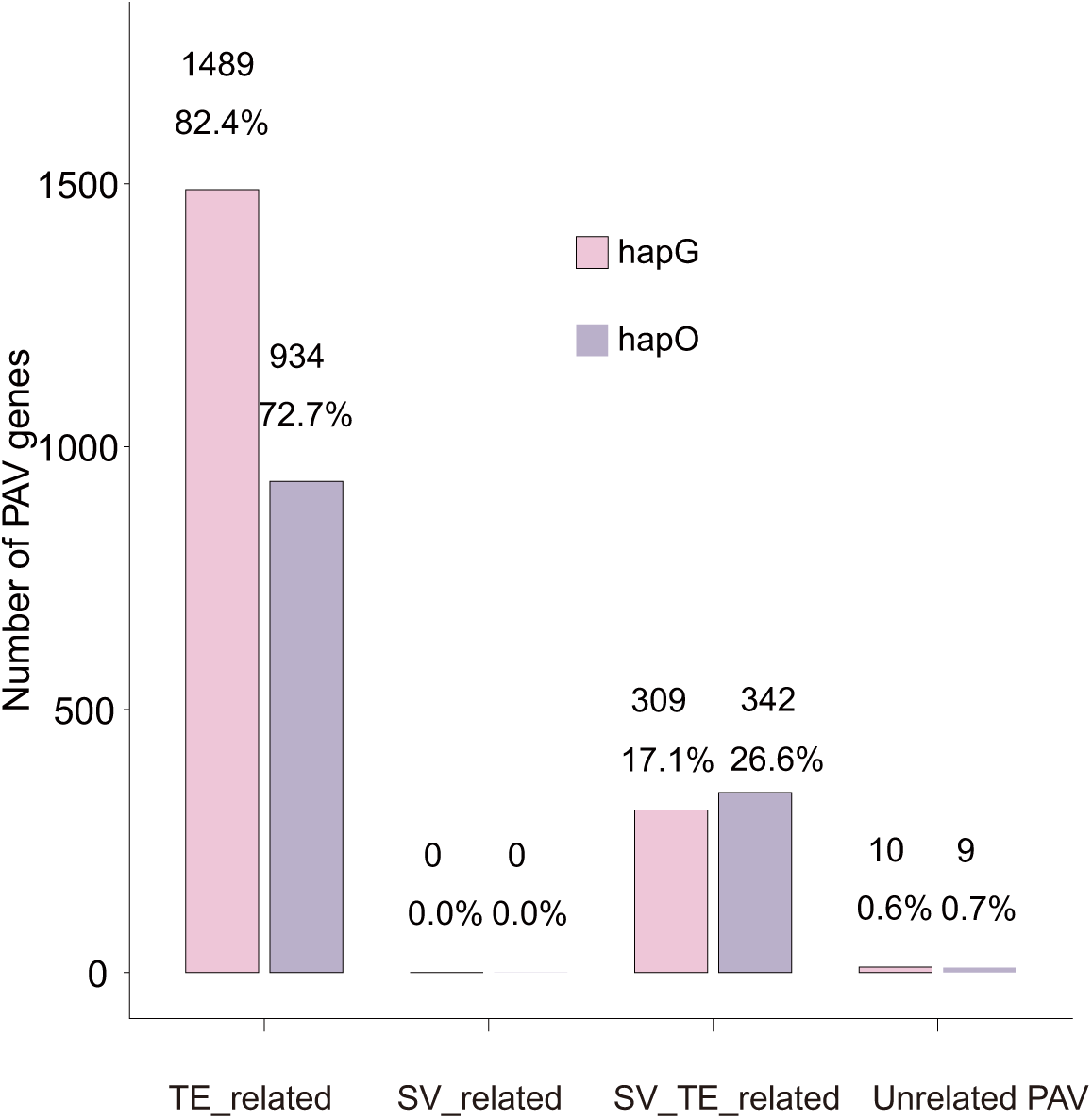
Contribution of transposable elements and structural variations to haplotype-specific PAV genes. Numbers and proportions of HapG- and HapO-specific PAV genes classified according to their association with transposable elements (TEs) and structural variations (SVs). In HapG, 1,489 PAV genes (82.4%) were TE-related, 309 (17.1%) were associated with both TEs and SVs, and 10 (0.6%) were unrelated to either category. In HapO, 934 PAV genes (72.7%) were TE-related, 342 (26.6%) were associated with both TEs and SVs, and 9 (0.7%) were unrelated. No exclusively SV-related PAV genes were identified in either haplotype.

**Figure S6.**
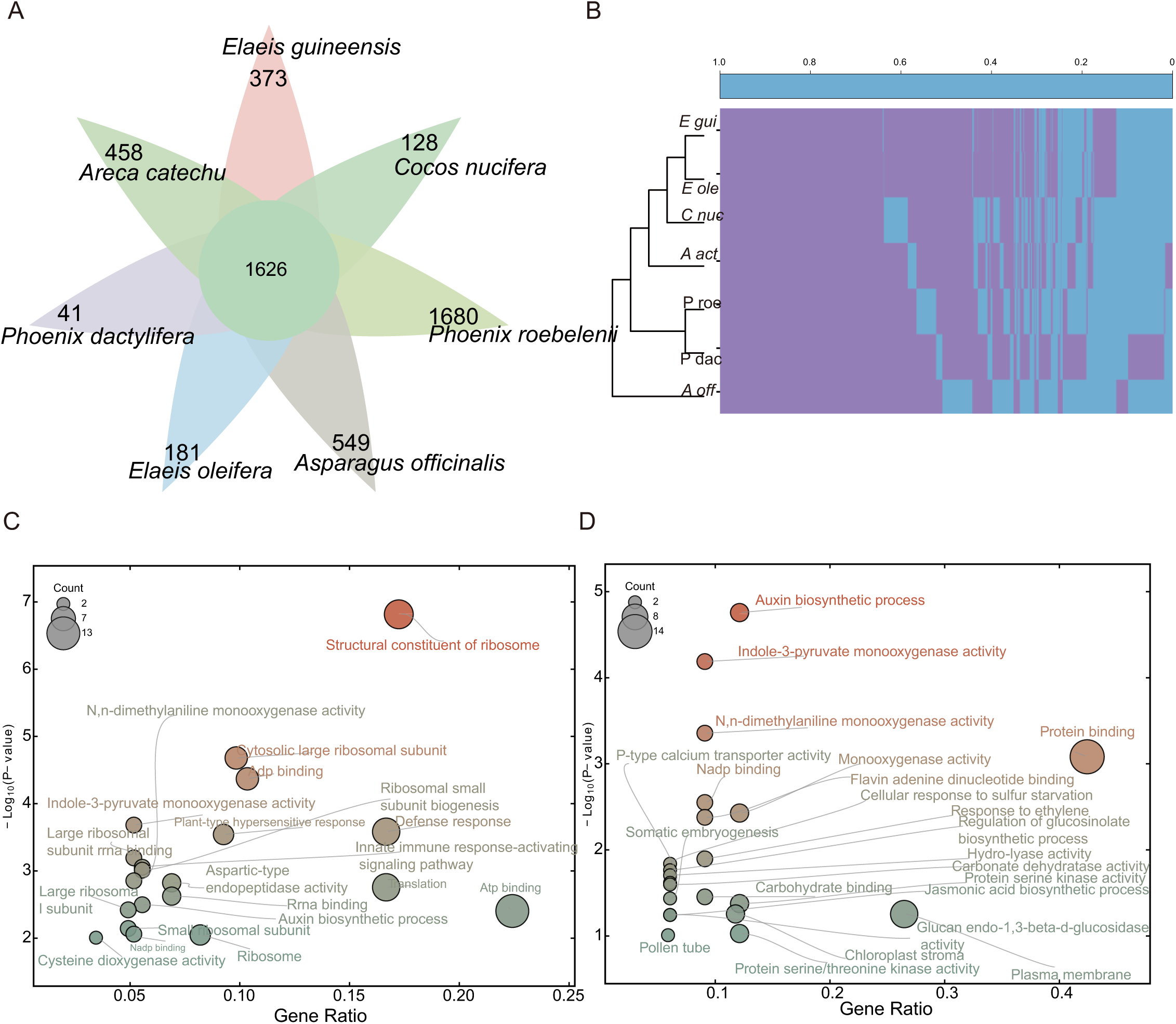
Comparative orthogroup analysis among seven monocot genomes. (A) Distribution of shared and species-specific orthogroups among *Elaeis guineensis*, *E. oleifera*, *Cocos nucifera*, *Areca catechu*, *Phoenix roebelenii*, *P. dactylifera*, and *Asparagus officinalis*. A total of 1,626 single-copy orthogroups were shared among all seven genomes; numbers at the tips indicate species-specific orthogroups. (B) Hierarchical clustering of orthogroup presence-absence patterns across the seven genomes. (C, D) Gene Ontology (GO) enrichment analyses of species-specific genes in *E. guineensis* and *E. oleifera*, respectively. Bubble size represents gene count, the x-axis represents gene ratio, and the y-axis represents -log10(P-value).

**Figure S7.**
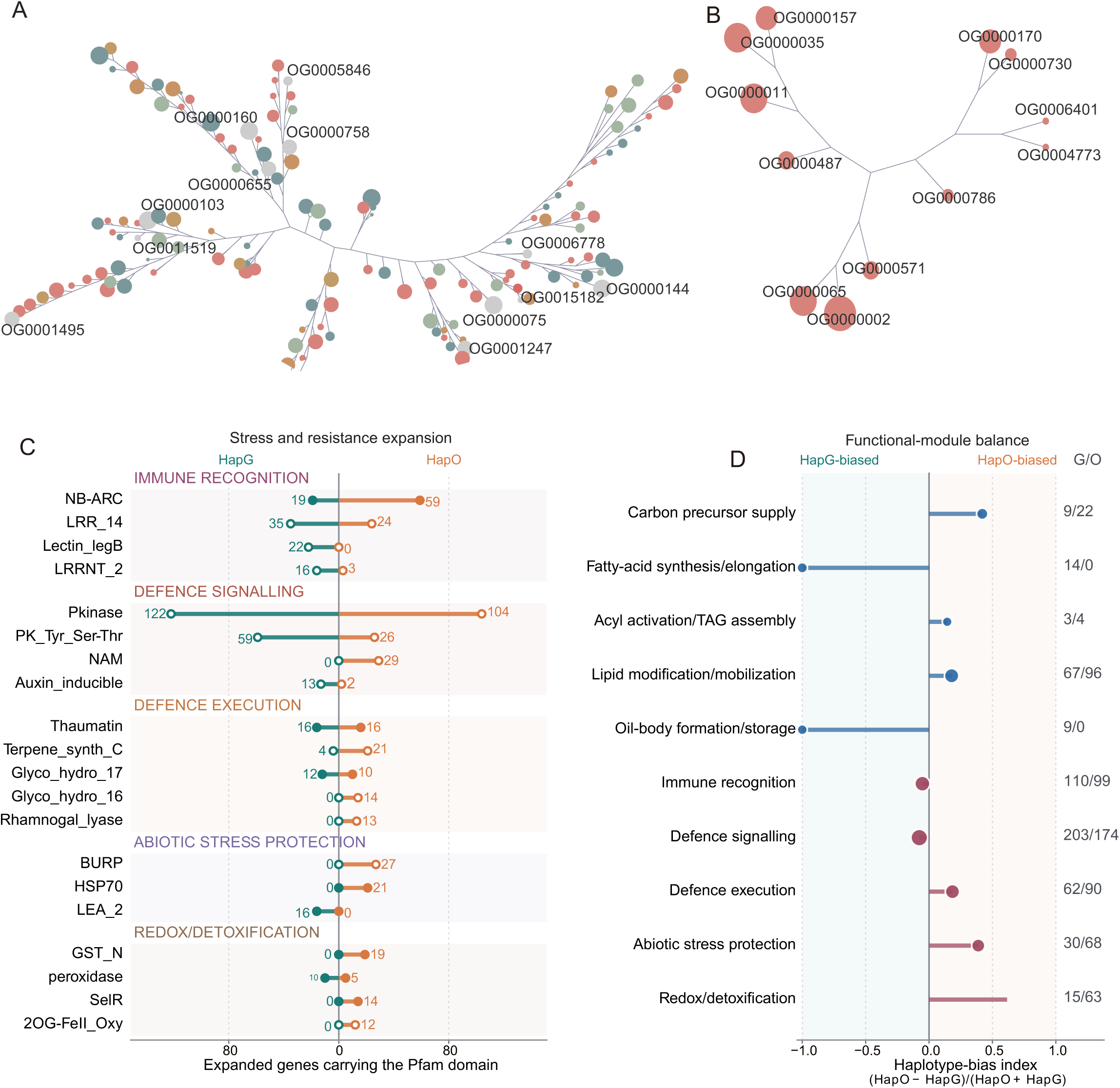
Functional characteristics of expanded gene families in HapG and HapO. (A) Phylogenetic tree of representative genes from selected HapG-expanded orthogroups in which *E. guineensis* showed the highest gene copy number among the analyzed species. (B) Phylogenetic tree of representative genes from selected HapO-expanded orthogroups in which *E. oleifera* showed the highest gene copy number among the analyzed species. (C) Comparison of expanded genes carrying representative Pfam domains between HapG and HapO. Domains are grouped into immune recognition, defense signaling, defense execution, abiotic stress protection, and redox/detoxification categories; numbers indicate expanded-gene counts in each haplotype. (D) Haplotype bias of functional modules related to lipid metabolism and stress responses. The haplotype-bias index was calculated as (HapO - HapG)/(HapO + HapG), with negative and positive values indicating HapG- and HapO-biased expansion, respectively. Numbers on the right indicate HapG/HapO gene counts for each functional module.

**Figure S8.**
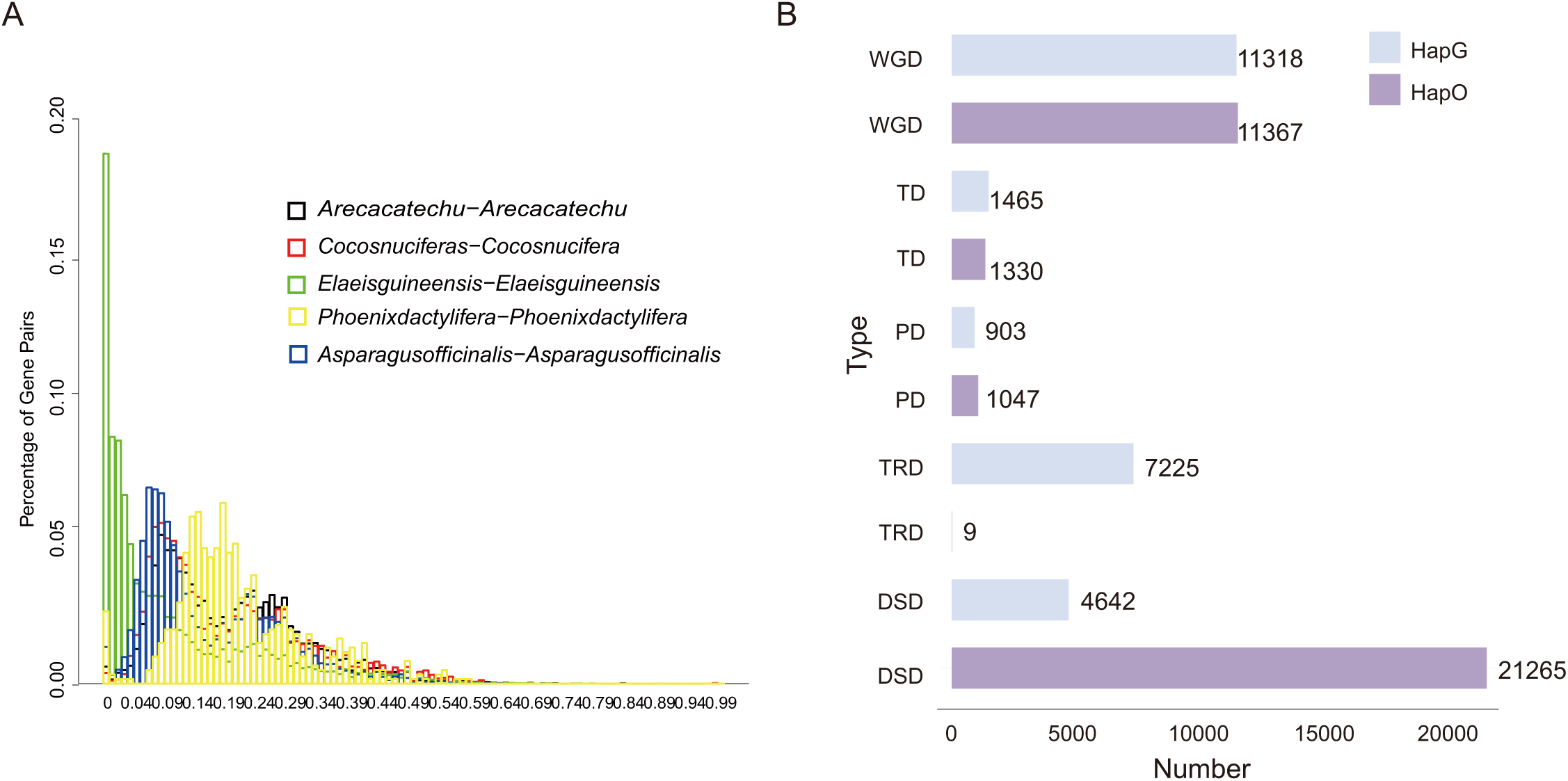
Synonymous-substitution distributions and gene-duplication modes. (A) Distribution of synonymous substitution rates among paralogous gene pairs in *Areca catechu*, *Cocos nucifera*, *Elaeis guineensis*, *Phoenix dactylifera*, and *Asparagus officinalis*. The y-axis represents the proportion of gene pairs within each interval. (B) Numbers of duplicated genes assigned to whole-genome or segmental duplication (WGD), tandem duplication (TD), proximal duplication (PD), transposed duplication (TRD), and dispersed duplication (DSD) in HapG and HapO.

**Figure S9.**
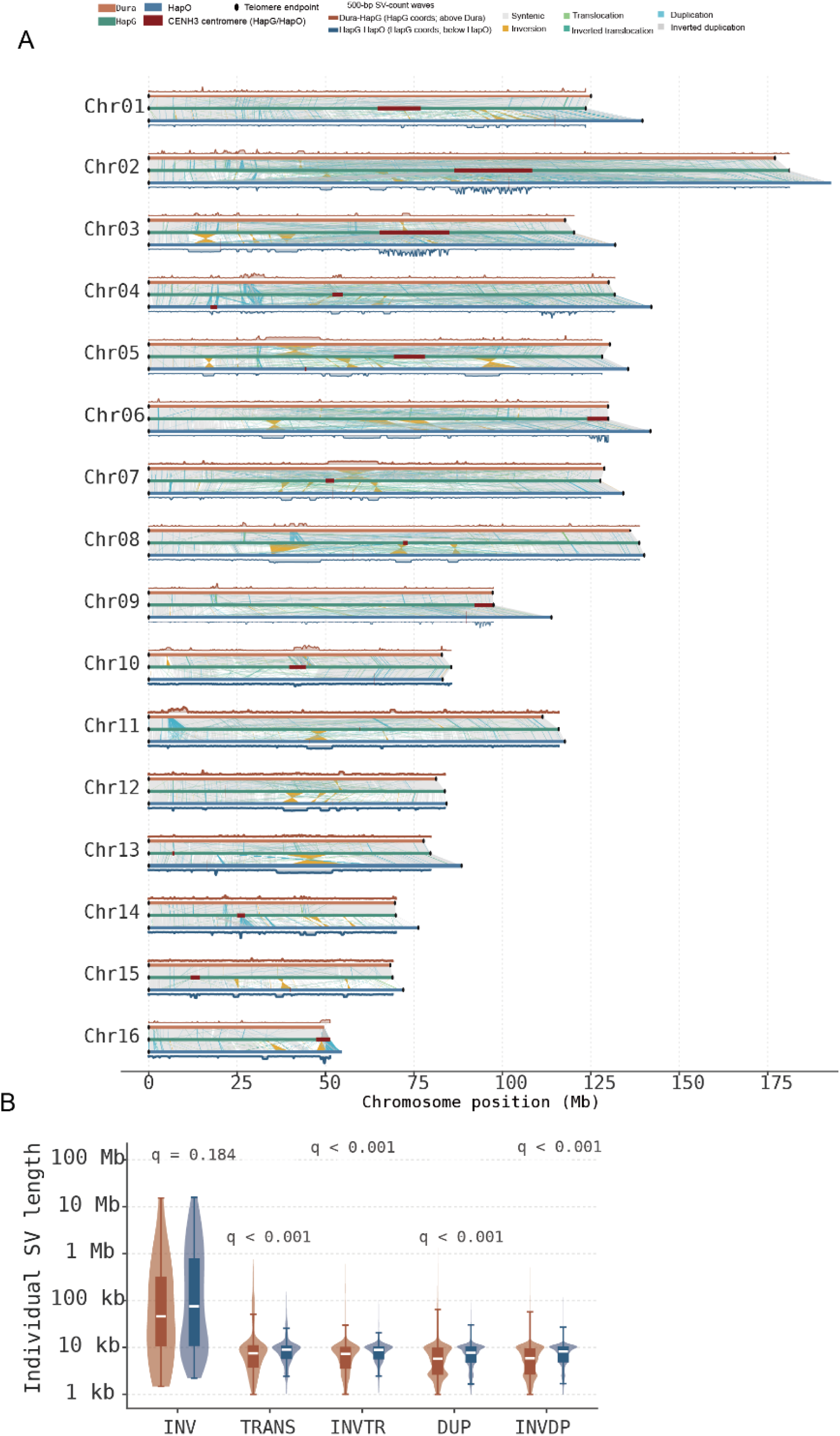
Genome-wide structural variation among Dura, HapG, and HapO. (A) Chromosome-scale synteny and major SV classes among Dura, HapG, and HapO, with CENH3-defined centromeres, telomeres, and 500-bp SV-count profiles indicated. (B) Size distributions of major SV classes in the Dura–HapG and HapG–HapO comparisons; adjusted *q*-values are shown for pairwise comparisons.

**Figure S10.**
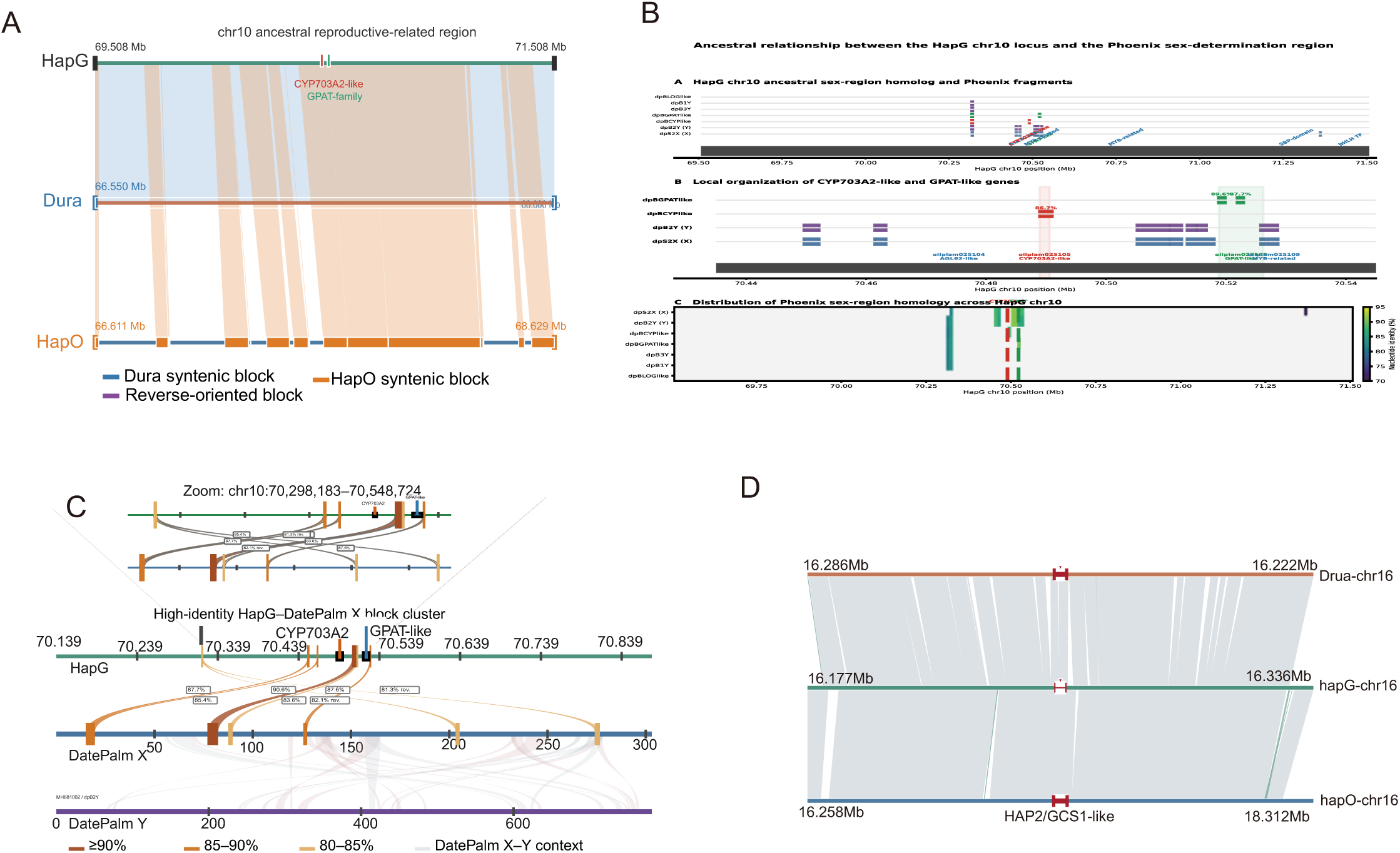
Conservation of reproductive-related genomic regions among oil palm genomes and date palm. The composite comparison summarizes conservation of reproductive-associated loci at chromosomes 10 and 16. The chromosome 10 analyses show the ancestral reproductive-related region in HapG and its syntenic relationships with Dura and HapO, together with *CYP703A2-like* and GPAT-family genes; Dura-syntenic, HapO-syntenic, and reverse-oriented blocks are distinguished. Comparisons with Phoenix sex-region fragments show high-identity HapG-DatePalm X blocks and the local organization of *CYP703A2*/*GPAT-like* genes, including a zoomed HapG interval at chr10:70,298,183-70,548,724. Alignment blocks are grouped by sequence identity (≥90%, 85-90%, and 80-85%), and the DatePalm X-Y genomic context is shown. The chromosome 16 comparison shows local synteny surrounding the *HAP2/GCS1-like* locus among Dura, HapG, and HapO.

**Figure S11.**
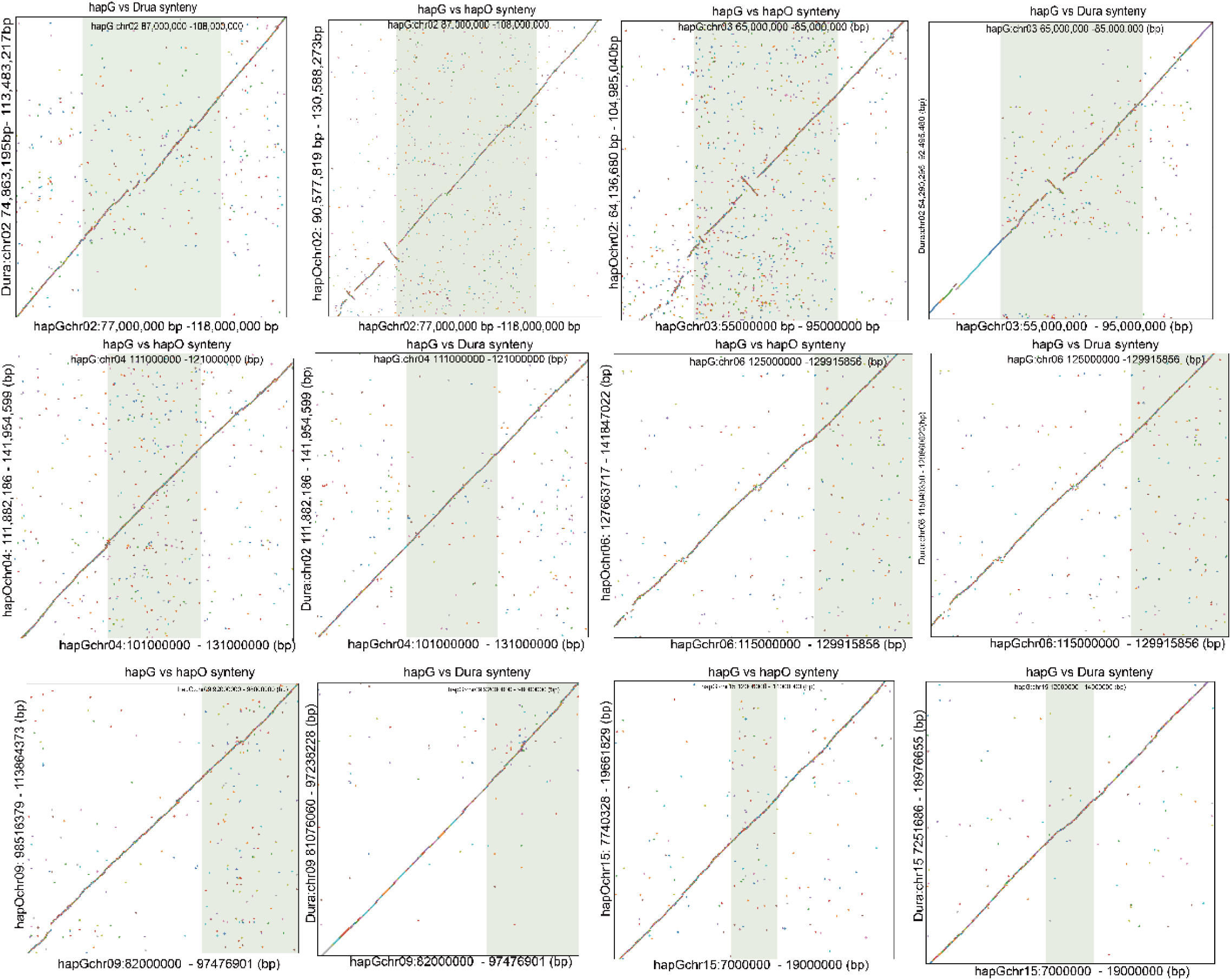
Pairwise synteny surrounding the six candidate introgressed regions in HapG. Dot-plot comparisons of the six candidate introgressed regions in HapG and their corresponding homologous regions in HapO and Dura on chromosomes 02, 03, 04, 06, 09, and 15. For each candidate region, HapG was compared separately with HapO and Dura. Green-shaded areas indicate the candidate introgressed intervals in HapG: chr02:87–108 Mb, chr03:65–85 Mb, chr04:111–121 Mb, chr06:125–129.916 Mb, chr09:92–98 Mb, and chr15:12–14 Mb. Flanking sequences are shown to illustrate local syntenic relationships and structural continuity

**Figure S12.**
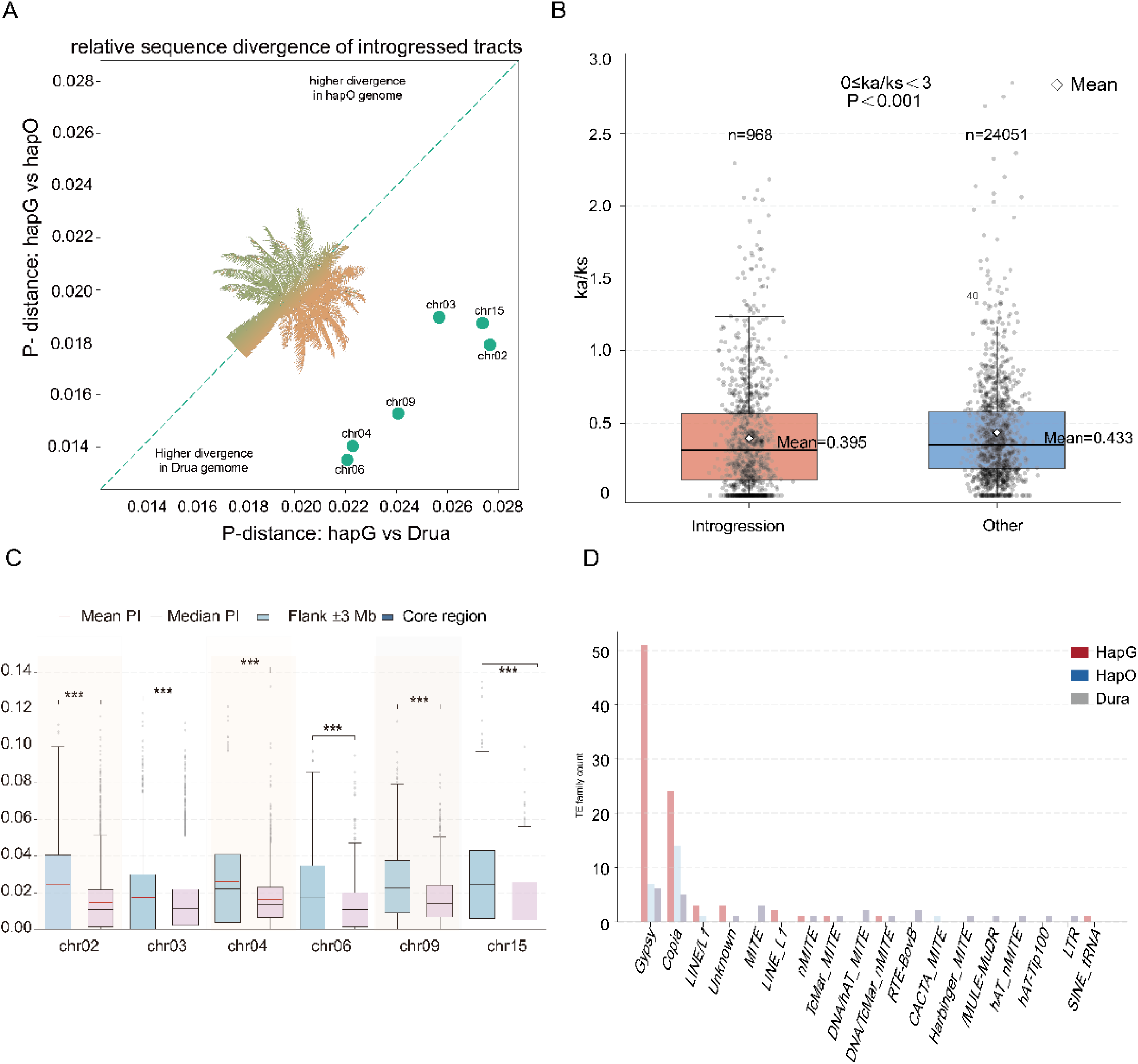
Evolutionary characteristics of the six candidate introgressed regions in HapG. (A) Relative sequence divergence of the six candidate introgressed regions. Pairwise p-distances between HapG and Dura are plotted against those between HapG and HapO; the dashed diagonal indicates equal divergence in the two comparisons. (B) Distribution of Ka/Ks ratios for genes within candidate introgressed regions and other genomic regions. Gene pairs with 0 ≤ Ka/Ks < 3 were included. Mean Ka/Ks values were 0.395 for introgressed regions (n = 968) and 0.433 for other genomic regions (n = 24,051; P < 0.001). (C) Nucleotide diversity (π) in the candidate introgressed cores and their ±3-Mb flanking regions on chromosomes 02, 03, 04, 06, 09, and 15. Mean and median π values are indicated; *** denotes P < 0.001. (D) Abundance of major transposable-element families within the ancient introgressed regions in HapG and their corresponding homologous regions in HapO and Dura.

**Figure S13.**
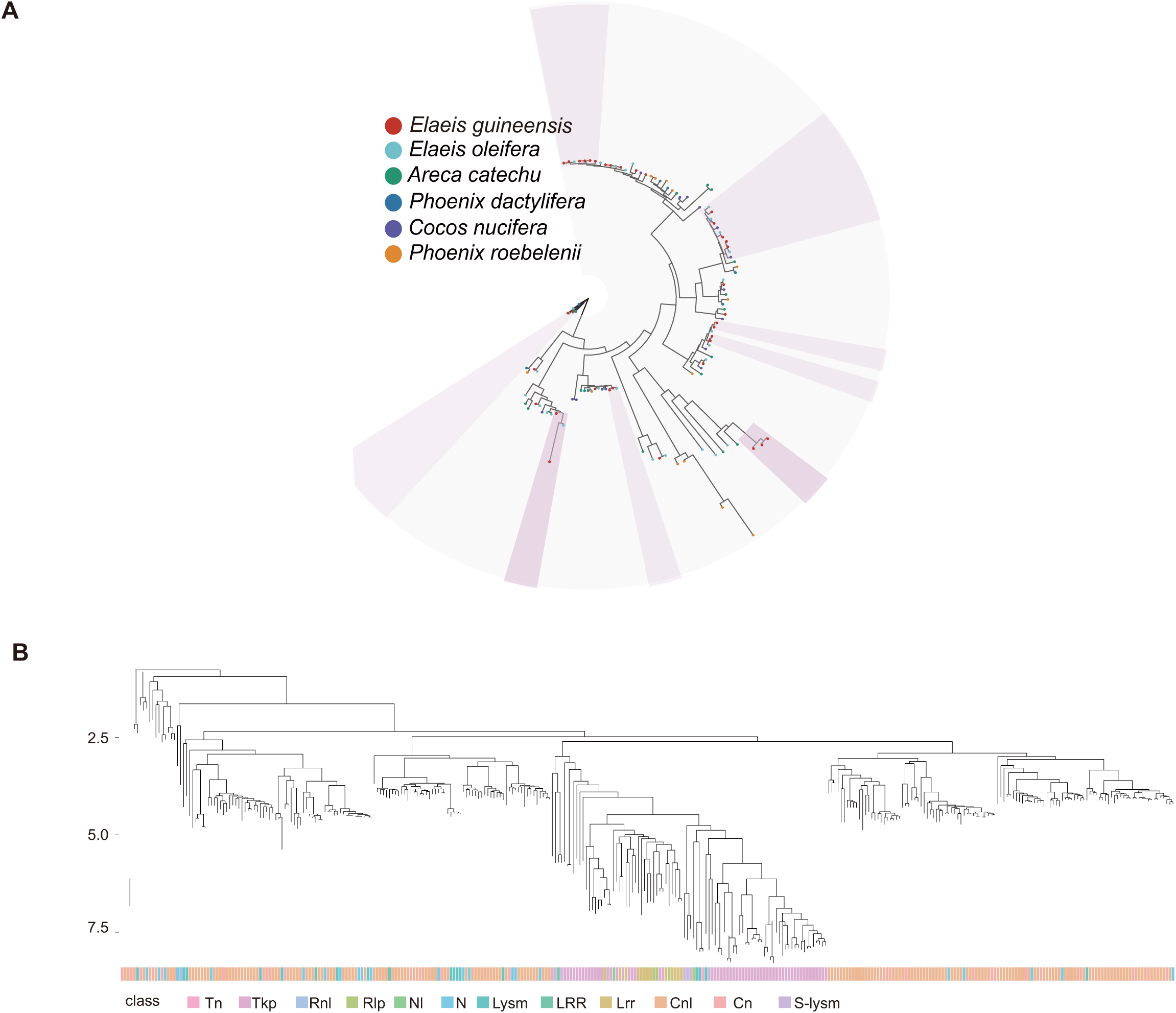
Phylogenetic relationships and classification of resistance-related gene families across palms. (A) Circular phylogenetic tree showing relationships among resistance-related proteins from *Elaeis guineensis*, *E. oleifera*, *Areca catechu*, *Phoenix dactylifera*, *Cocos nucifera*, and *P. roebelenii*. Terminal colors indicate species of origin. (B) Phylogenetic distribution of resistance-related classes, including Tn, Tkp, Rnl, Rlp, Nl, N, LysM, LRR, Lrr, Cnl, Cn, and S-LysM. Colored bars below the tree indicate the corresponding classes.

**Figure S14.**
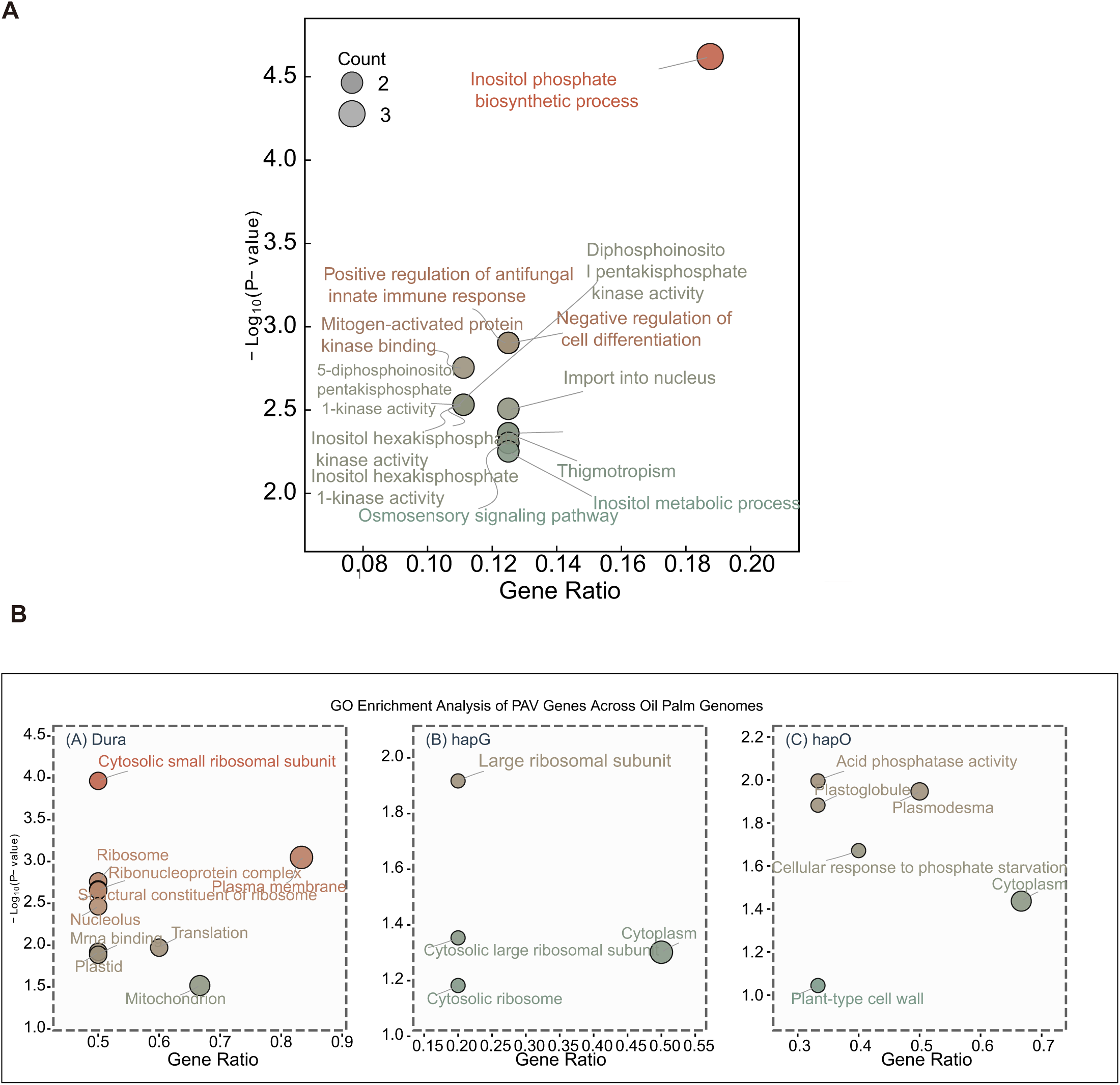
Functional enrichment associated with ancient introgressed and highly differentiated genomic regions. (A) GO enrichment profile of genes associated with highly differentiated HapG segments, highlighting terms related to inositol phosphate biosynthesis and metabolism, antifungal innate immune response, osmosensory signaling, thigmotropism, MAP kinase binding, and inositol phosphate kinase activities. (B) Comparative GO enrichment analysis of PAV genes within the ancient introgressed regions and their corresponding homologous regions in Dura, HapG, and HapO. The x-axis represents the gene ratio, the y-axis represents −log10(*P*-value), and bubble size indicates the number of genes associated with each GO term.

**Figure S15.**
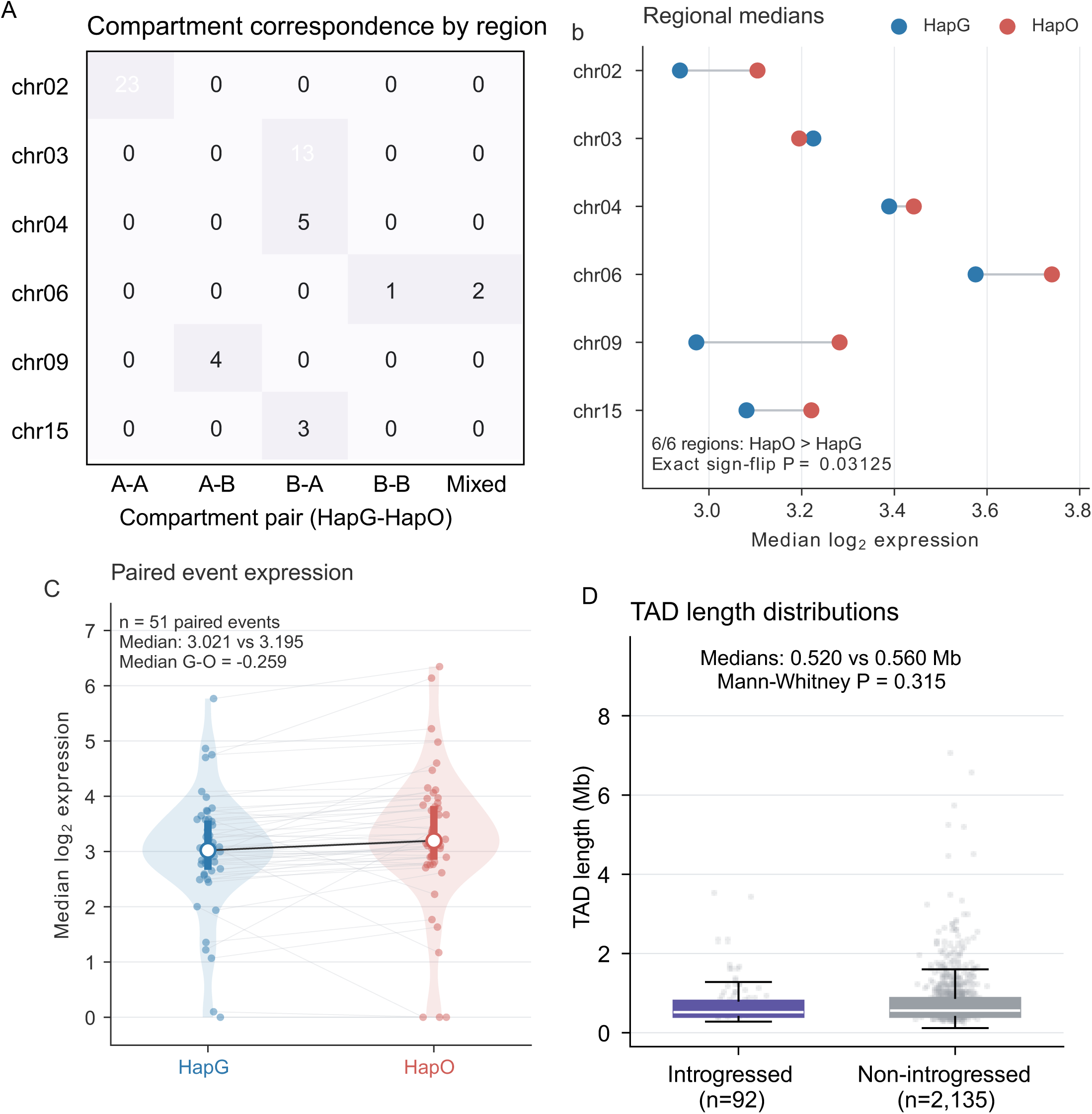
Chromatin organization and expression characteristics of candidate introgressed regions. (A) Correspondence of A/B chromatin compartments between HapG and HapO across the six candidate introgressed regions. Numbers indicate counts assigned to each HapG-HapO compartment combination (A-A, A-B, B-A, B-B, or mixed). (B) Median log2-transformed expression in HapG and HapO across the six candidate regions; HapO was higher in all six regions (exact sign-flip test, P = 0.03125). (C) Paired median expression for 51 matched events. Median log2 expression was 3.021 in HapG and 3.195 in HapO, with a median HapG-HapO difference of -0.259. (D) TAD-length distributions in introgressed (n = 92) and non-introgressed (n = 2,135) regions. Median TAD lengths were 0.520 and 0.560 Mb, respectively (two-sided Mann-Whitney U test, P = 0.315).

**Figure S16.**
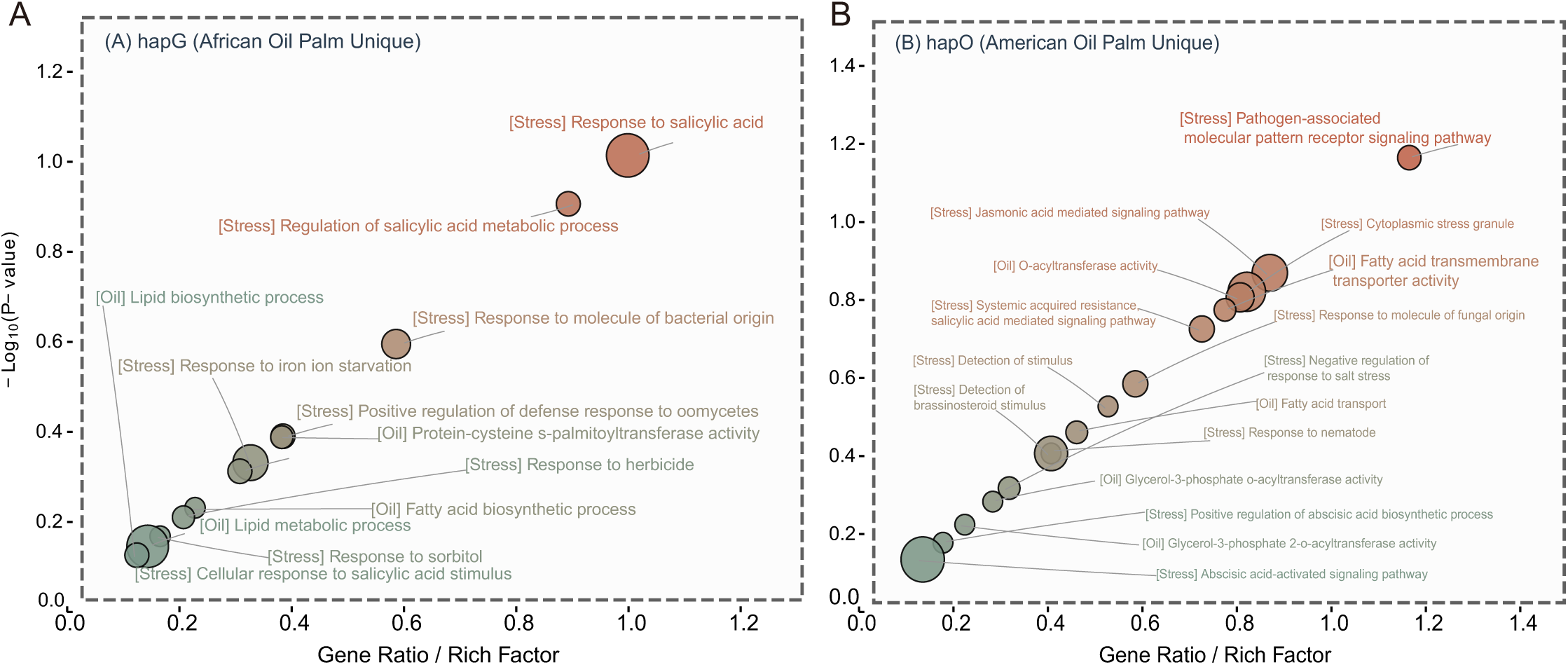
Functional enrichment of allelic genes with HapG- or HapO-dominant expression. (A) GO enrichment analysis of allelic genes showing HapG-dominant expression, highlighting functions related to lipid biosynthesis and metabolism as well as stress-response processes involving salicylic acid, bacterial-origin molecules, iron starvation, herbicides, and oomycetes. (B) GO enrichment analysis of allelic genes showing HapO-dominant expression, highlighting pathogen-associated molecular pattern receptor signaling, jasmonic acid- and salicylic acid-mediated responses, fungal and nematode responses, abscisic acid signaling, fatty-acid transport, and glycerol-3-phosphate acyltransferase activities. The x-axis represents the gene ratio/rich factor, and the y-axis represents −log10(*P*-value).

**Figure S17.**
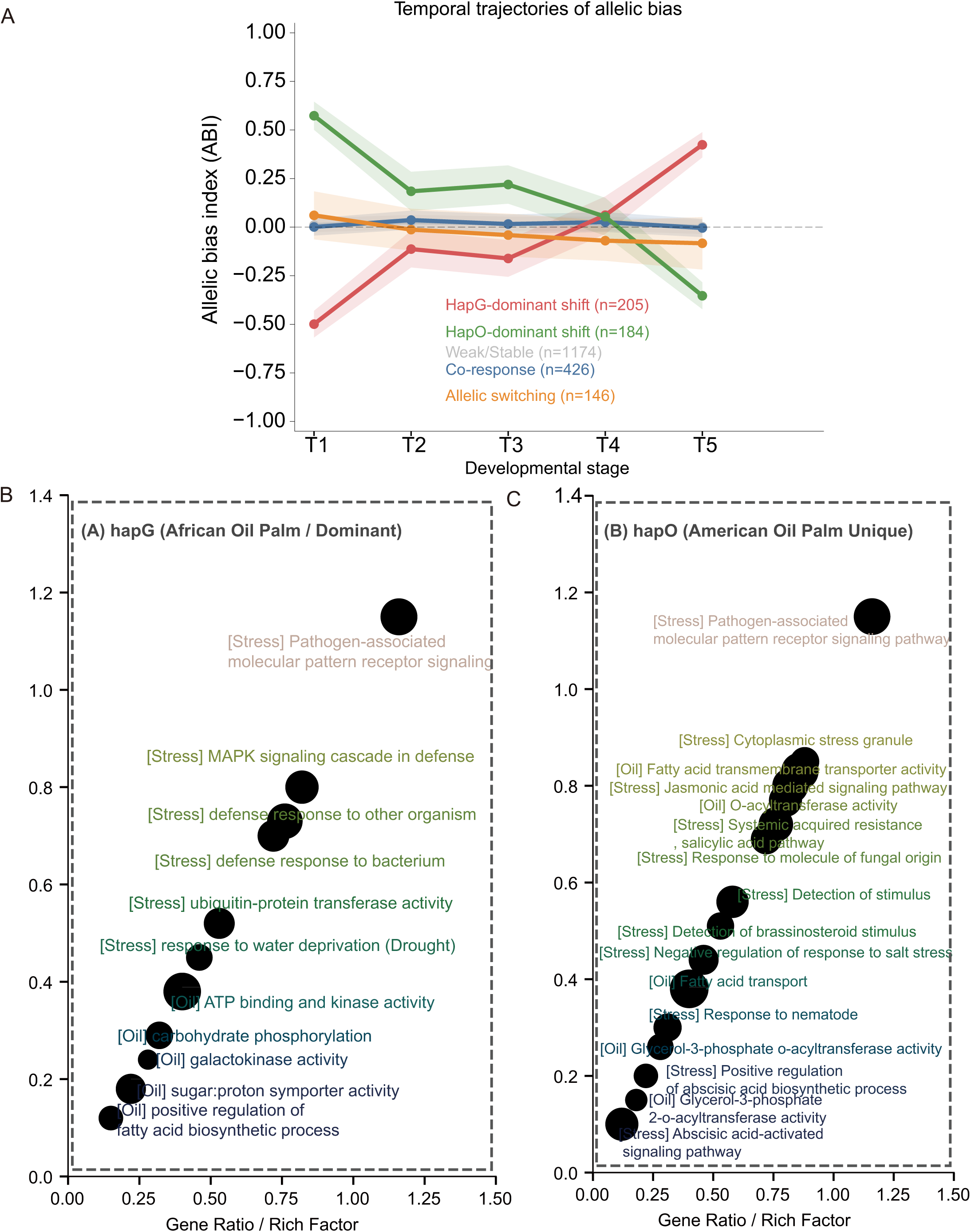
Allelic-bias dynamics and functional enrichment of loci with HapG- or HapO-biased expression. (A) Temporal trajectories of the allelic bias index (ABI) across five developmental stages (T1–T5), showing distinct patterns of allelic expression dynamics among paired HapG–HapO loci. (B) GO enrichment analysis of loci showing HapG-biased expression, highlighting functions related to fatty-acid biosynthesis, carbohydrate metabolism, kinase activity, drought response, defense responses, MAPK signaling, and pathogen-associated molecular pattern receptor signaling. (C) GO enrichment analysis of loci showing HapO-biased expression, highlighting functions related to glycerol-3-phosphate acyltransferase activity, fatty-acid transport, abscisic acid-, salicylic acid-, and jasmonic acid-mediated responses, salt-stress responses, and fungal and nematode responses. For panels B and C, enrichment analyses were performed using genes corresponding to HapG- or HapO-biased allelic loci, respectively. The x-axis represents the gene ratio/rich factor, and the y-axis represents −log10(*P*-value).

